# Osteocytes regulate organismal senescence of bone and bone marrow

**DOI:** 10.1101/2022.07.17.500353

**Authors:** Peng Ding, Chuan Gao, Youshui Gao, Delin Liu, Hao Li, Jun Xu, Xiaoyi Chen, Yigang Huang, Changqing Zhang, Minghao Zheng, Junjie Gao

**Affiliations:** Department of Orthopaedics, Shanghai Jiao Tong University Affiliated Sixth People’s Hospital, Shanghai, 200233, China; Centre for Orthopaedic Translational Research, Medical School, University of Western Australia, Nedlands, Western Australia 6009, Australia; Perron Institute for Neurological and Translational Science, Nedlands, Western Australia 6009, Australia; Ningbo Institute of Life and Health Industry, University of Chinese Academy of Sciences; Institute of Microsurgery on Extremities, Shanghai Jiao Tong University Affiliated Sixth People’s Hospital, Shanghai, 200233, China

**Keywords:** Osteocytes, SASP, osteogenesis, osteoclastogenesis, myelopoiesis

## Abstract

The skeletal system contains a series of sophisticated cellular lineages arisen from the mesenchymal stem cells (MSC) and hematopoietic stem cells (HSC), that determine the homeostasis of bone and bone marrow. Here we reasoned that osteocyte may exert a function in regulation of these lineage cell specifications and tissue homeostasis. Using a mouse model of conditional deletion of osteocytes by the expression of diphtheria toxin subunit *α* (DTA) in dentin matrix protein 1 (DMP-1) positive osteocytes, we demonstrated that partial ablation of DMP-1positive osteocytes caused severe sarcopenia, osteoporosis and degenerative kyphosis, leading to shorter lifespan in these animals. Osteocyte reduction altered mesenchymal lineage commitment resulting in impairment of osteogenesis and induction of osteoclastogensis. Single cell RNA sequencing further revealed that hematopoietic lineage was mobilized towards myeloid lineage differentiation with expanded myeloid progenitors, neutrophils and monocytes, while the lymphopoiesis was impaired with reduced B cells in the osteocyte ablation mice. The acquisition of a senescence-associated secretory phenotype (SASP) in both osteoprogenic and myeloid lineage cells was the underlying cause. Together, we showed that osteocytes play critical roles in regulating of lineage cell specifications in bone and bone marrow through mediation of organismal senescence.

## Introduction

The skeletal system is an elaborate organ mainly containing bone, bone marrow and other connective tissues, whose function includes movement, support, hematopoiesis, immune responses and endocrine regulation(Karsenty and Ferron 2012; Katsnelson 2010; Quarles 2011). The skeletal system hosts at least more than 12 types of cell lineage differentiations arisen from the hematopoietic stem cells (HSC) and mesenchymal stem cells (MSC)(Mendez-Ferrer et al. 2010). During hematopoiesis, HSCs give rise to lymphoid and myeloid lineage cells including B cell, neutrophil and monocytes as well as osteoclasts. Meanwhile, MSCs differentiate into osteoblastic lineage cells, bone marrow adipocytes and form fibroconnective tissues. The sophisticated processes of differentiation and interaction of these cell lineages are critical not only to skeletal development, but also to the integrity of hematopoietic, immune and endocrine systems(Mendez-Ferrer, et al. 2010; Le, Andreeff, and Battula 2018; Yu and Scadden 2016). During aging, these cell lineage commitments change rigorously and cause imbalance between myeloid-lymphoid hematopoiesis and adipo-osteogenic differentiation (Chen et al. 2016; Sinha et al. 2022), which lead to the increased myelopoiesis and adipogenesis as opposed to lymphopoiesis and osteogenesis. While the complex communications between theses cell lineages have been documented, it is still unclear what determine these cell lineages to survive and how their cell fates are maintained during development and aging. It has been speculated that cellular senescence, characterized by cell proliferation arrest, altered metabolism and apoptosis resistance(Gorgoulis et al. 2019; Tchkonia et al. 2013), may be responsible for the regulation of lineage cell fates. However, the precise role in aging and age-related diseases remain unclear.

Osteocytes, as the long living terminally differentiated cells and the most abundant cells within the bone matrix(Tresguerres et al. 2020), play vital roles in maintaining the skeletal homeostasis. Apart from mechanical transduction(Long 2011; Sato et al. 2020), osteocytes have been shown to regulate bone formation, bone resorption, bone marrow hematopoiesis(Asada et al. 2013; Azab et al. 2020; Fulzele et al. 2013; Xiao et al. 2021) and generate endocrine signals to mediate function of other organs(Razzaque 2009; Fulzele et al. 2017; Cain et al. 2012). Here we hypothesize that osteocytes, may exert another important role in regulation of lineage cell fate specifications, and harmonization of bone and bone marrow through mediation of organismal senescence. Using a mouse model of conditional deletion of osteocytes by the expression of diphtheria toxin subunit *α* (DTA) in dentin matrix protein 1 (DMP-1) positive osteocytes, we showed that osteocytes regulated organismal senescence of bone and bone marrow resulted in skeletal premature aging including severe sarcopenia, osteoporosis and kyphosis. Deletion of DMP-1 positive osteocytes in mouse impaired osteogenesis, increased osteoclastogenesis and myelopoiesis. HSCs were mobilized towards myeloid lineage differentiation with expanded myeloid progenitors, neutrophils and monocytes, while the lymphopoiesis was impaired with reduced B cells. Together, we demonstrated that osteocyte played a critical role in regulation of the HSC and MSC lineage cell differentiations by modification of organismal senescence.

## Results

### Mice with less osteocytes have severe osteoporosis, kyphosis, sarcopenia and shorter lifespan

To delineate the role of osteocyte in skeletal tissue development and maturation, we established a mouse model based on diphtheria toxin subunit α-mediated cell knockout using the promoter of DMP-1(Breitman et al. 1990). The latter is a protein highly expressed in late stage osteocytes but has been shown not to be essential for early skeletal development(Feng et al. 2003). The results showed that complete ablation of DMP-1 positive osteocytes (osteocyte^DMP-1^) in DMP-1^cre^ DTA^fl/fl^ mice (DTA^ho^) caused lethality of mice before birth. This has led us to investigate the impact of partial ablation of osteocytes using DMP-1^cre^ DTA^fl/+^ mice (DTA^het^). As shown in Figure 1A and B, DTA^het^ mice had more empty lacunae without the presence of osteocytes within cortical and trabecular bone matrix as compared to WT mice. Further, reduced dendrites were also observed in residual osteocytes of DTA^het^ mice (Figure 1C and D), indicating that the impairment of osteocyte network. Interestingly, Alizarin red/Alcian blue staining of whole mount skeleton at E19.0 showed no apparent differences of craniofacial, long bones or spines between WT and DTA^het^ mice (Figure 2 - figure supplement 1). Together, these results indicated that although there was partial ablation of osteocyte^DMP-1^ in DTA^het^ mice, the embryonic development of skeletal tissue appeared to be normal.

**Figure 1.**
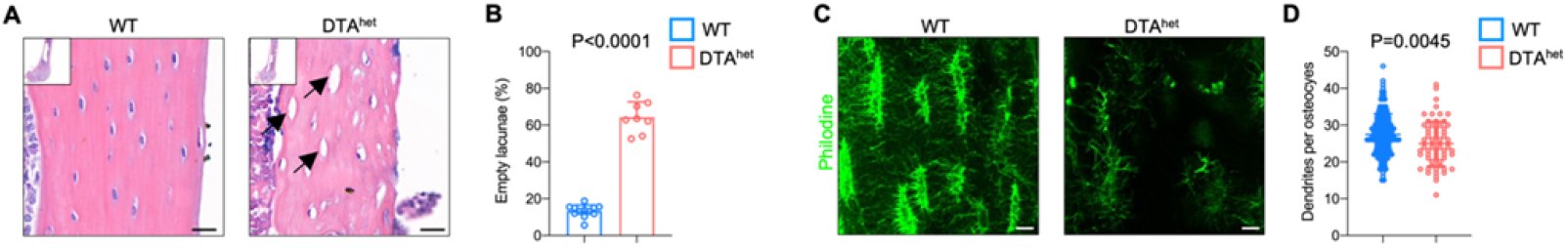
DTA^het^ mice display partial osteocyte ablation. (**A-B**) Hematoxylin-eosin staining of WT and DTA^het^ mice femur at 4 weeks (**A**) and quantification of the ratio of empty lacunae (arrows) (**B**) (n=8-12 per group), indicating reduced osteocyte number in DTA^het^ mice. Scare bar, 20 μm. (**C-D**) Immunofluorescence staining of femoral cortical bone of 4-week-old WT and DTA^het^ mice (**C**) and quantification of dendrites per osteocyte based on the images (**D**) (n=152 osteocytes in WT group and n=64 osteocytes in DTA^het^ group). Scare bar, 20 μm. Error bar represents the standard deviation.

Next, we investigated if reduction of osteocyte^DMP-1^ in DTA^het^ mice had an impact of postnatal maturation of bone tissue. Micro-computed tomography (μCT) examination of the appendicular skeleton revealed a significant decrease in femur bone mineral density (BMD), bone volume fraction (BV/TV), trabecular number (Tb.N) and trabecular thickness (Tb.Th), as well as greater trabecular separation (Tb.Sp) in DTA^het^ mice as compared to those in WT mice at 4 weeks (Figure 2A and B). Moreover, ablation of osteocytes also led to cortical bone loss with decreased cortical thickness (Ct.Th) and increased cortical porosity (Ct.Po) (Figure 2A and C). At 13 weeks, DTA^het^ mice exhibited more bone loss in both trabecular and cortical bone compared to those in WT mice (Figure 2D-G). The progressive bone loss was observed through the life of DTA^het^ mice. The phenotype observed is unique and gender insensitive (Figure 2 - figure supplement 2A-C). Similarly, μCT observation of axial skeleton also revealed the significant bone loss in vertebral bodies (Fig 2H and I, Figure 2 - figure supplement 2D and E). Furthermore, there was no increase of bone mass of vertebral bodies from 4 to13 weeks in DTA^het^ mice (Figure 2H and I), suggesting retardation of vertebral body maturation. At 13 weeks, obvious kyphosis occurred in DTA^het^ mice (Figure 2L) due to serve osteoporosis and vertebral body compression. Whole-body μCT scan revealed that there was giant increase of thoracic and lumbar curvature of DTA^het^ mice (Figure 2M). At the age of 20 weeks almost all of DTA^het^ mice developed severe kyphosis (Figure 2N). In consistent with development of kyphosis, gait analysis revealed that DTA^het^ mice at 4 weeks have abnormal steps when running (Figure 2 - figure supplement 3A and B). The font and hind stride length were much shorter in DTA^het^ mice (Figure 2 - figure supplement 3C). Also, the swing speed of DTA^het^ mice was much slower than WT mice (Figure 2 - figure supplement 3D, Movie supplement 1-6).

**Figure 2.**
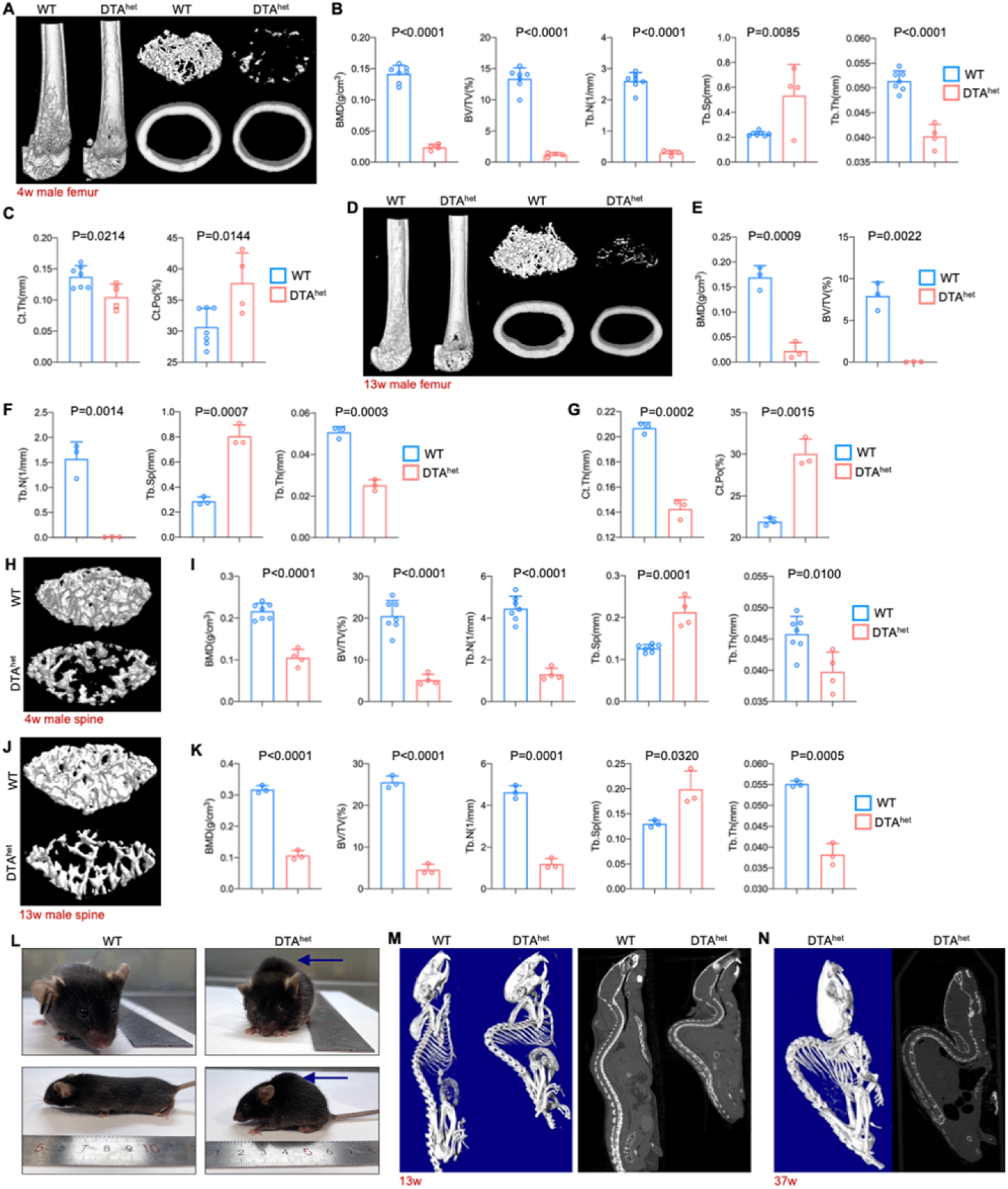
Osteocyte ablation induces severe osteoporosis and kyphosis. (**A-C**) Representative μCT reconstructive images of male WT and DTA^het^ mice femur at 4 weeks (**A**) and trabecular microstructural parameters (bone mineral density, BMD; bone volume fraction, BV/TV; trabecular number, Tb.N; trabecular separation, Tb.Sp; and trabecular thickness, Tb.Th); (**B**) and cortical microstructural parameters (cortical thickness, Ct.Th; and cortical porosity, Ct.Po) (**C**) derived from μCT analysis (n=4-7 per group). (**D-G**) Representative μCT reconstructive images of male WT and DTA^het^ mice femur at 13 weeks (**D**) and trabecular microstructural parameters (BMD, BV/TV, Tb.N, Tb.Sp and Tb.Th) (**E-F**) and cortical microstructural parameters (Ct.Th and Ct.Po) (**G**) derived from μCT analysis (n=3 per group), demonstrating severe bone loss in DTA^het^ mice. (**H-I**) Representative μCT reconstructive images of male WT and DTA^het^ mice third lumbar at 4 weeks (**H**) and trabecular microstructural parameters (BMD, BV/TV, Tb.N, Tb.Sp and Tb.Th) (**I**) derived from μCT analysis. (**J-K**) Representative μCT reconstructive images of male WT and DTA^het^ mice third lumbar at 13 weeks (**J**) and trabecular microstructural parameters (BMD, BV/TV, Tb.N, Tb.Sp and Tb.Th) (**K**) derived from μCT analysis, showing vertebral body bone loss in the spine of DTA^het^ mice. (**L**) Gross images of male WT and DTA^het^ mice at 13 weeks. (**M**) Representative whole-body μCT reconstructive and sagittal images of male WT and DTA^het^ mice at 13 weeks. (**N**) Representative whole-body μCT reconstructive and sagittal images of male DTA^het^ mice at 37 weeks, noting that severe kyphosis occurred in DTA^het^ mice. Error bar represents the standard deviation.

Whole body examination of DTA^het^ mice revealed there was a continual body weight loss and muscle weight loss (Figure 3A, B and C) from 4 weeks. Histology examination of gastrocnemius muscles revealed focal muscle atrophy with mild inflammation at 4 weeks (Figure 3D and E). No muscle fibrosis was observed. Many myonuclei were mispositioned and became centralized as contrast to those in WT mice. At 13 weeks, there was continual muscle atrophy, rimmed vacuoles and inclusion bodies were seen within the muscle fibers (Figure 3F-G). Together these results suggested that DTA^het^ mice had systemic muscle atrophy and sarcopenia. Subsequently, the average lifespan of DTA^het^ mice was about 20-40 weeks, which was much shorter than WT mice (Figure 3H). Together, these data demonstrated that osteocytes ablation caused severe osteoporosis and kyphosis, as well as sarcopenia. These premature aging phenotypes have resulted in shortened lifespan.

**Figure 3.**
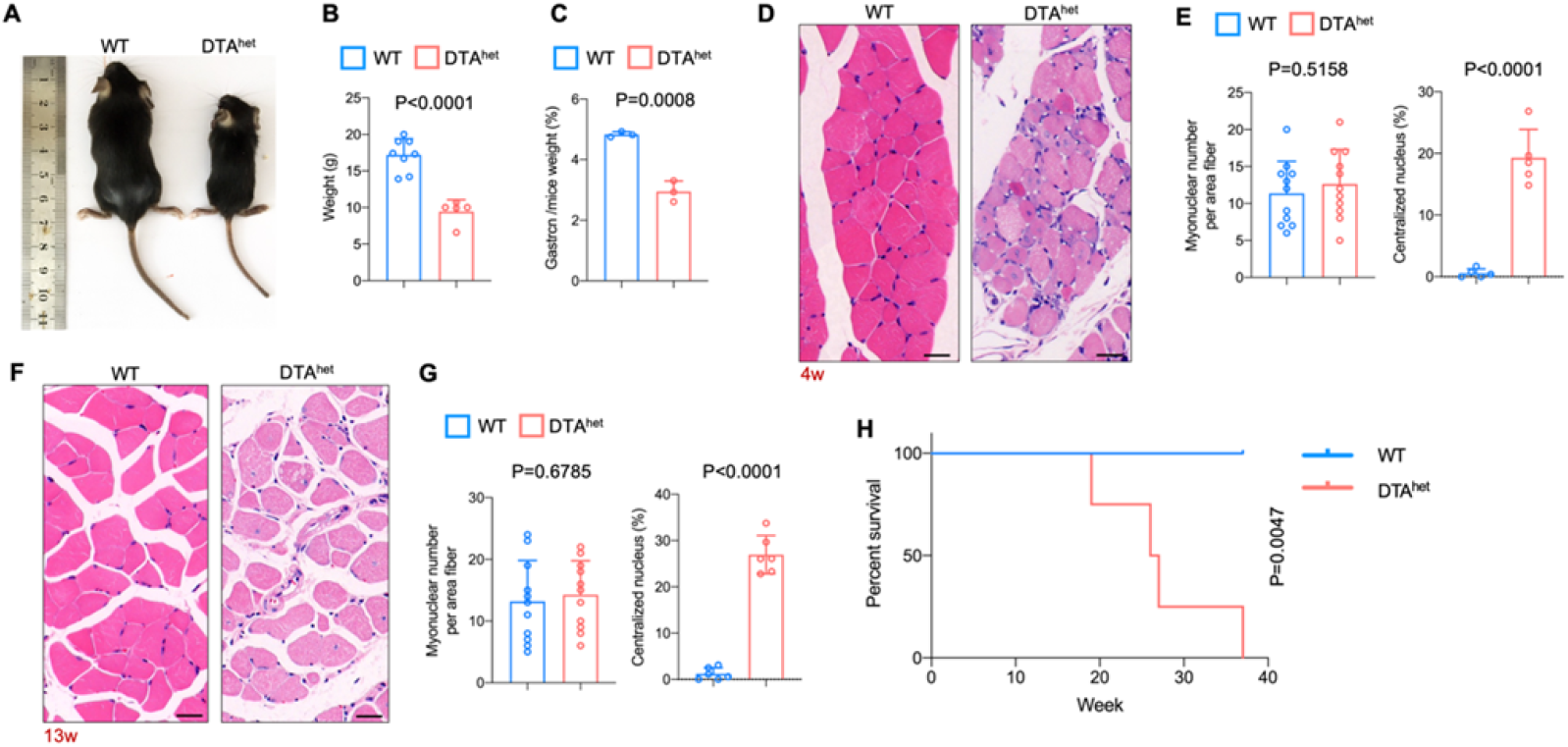
Osteocyte ablation leads to severe sarcopenia and shorter lifespan. (**A-B**) Gross images (**A**) and weight (**B**) of male WT and DTA^het^ mice at 4 weeks (n=5-8 per group). (**C**) The ratio of gastrocnemius muscle weight male WT and DTA^het^ mice at 4 weeks (n=3 per group). (**D-E**) Hematoxylin-eosin staining of WT and DTA^het^ mice gastrocnemius muscle at 4 weeks (**D**) and quantification of myonuclei per area fiber (n=11 per group) and centralized nucleus per field (**E**) (n=5 per group). Scale bar, 20 μm. Showing focal muscle atrophy, increased centralized myonuclei and mild inflammation in DTA^het^ mice. (**F-G**) Hematoxylin-eosin staining of WT and DTA^het^ mice gastrocnemius muscle at 13 weeks (**F**) and quantification of myonuclei per area fiber (n=11 per group) and centralized nucleus per field (**G**) (n=6 per group). Noting muscle atrophy, rimmed vacuoles and inclusion bodies within the muscle fibers in DTA^het^ mice. Scale bar, 20 μm. (**H**) Kaplan-Meier survival curve of WT and DTA^het^ mice (n=4-5 per group), showing that DTA^het^ mice had shorter lifespan than that of wild type. Error bar represents the standard deviation.

### Ablation of osteocytes alters mesenchymal lineage commitment and promoted osteoclastogensis

To explore the potential mechanism on why reduction of osteocytes has caused severe osteoporosis and kyphosis, RNA sequencing was performed on whole bone with bone marrow flushed out from DTA^het^ and WT mice at 4 weeks. Selected skeleton related gene ontology (GO) analysis revealed that downregulated genes by osteocyte ablation were enriched in ossification, osteoblast differentiation, positive regulation of osteoblast differentiation, endochondral ossification and bone morphogenesis (Figure 4 - figure supplement 1A and Supplementary file table 1). Heatmap of significantly differentiated genes (fold change > 2.0-fold, WT average FPKM > 10, FDR < 0.05) and subsequent RT-qPCR verified that genes that are critical for osteogenesis, including Alp, Ocn, Col1a1, Opn, Osx and Runx2, were affected by the ablation of osteocytes (Figure 4 - figure supplement 1B and C). In addition, numbers of osteoblasts and osteoid surface were remarkably reduced in DTA^het^ mice compared to WT (Figure 4A and B). Also, bone marrow fat accumulation in DTA^het^ mice was observed (Figure 4C and D). Together these results suggested that DTA^het^ mice displayed increased adipogenesis and decreased osteogenesis. To further evaluate the dynamics of bone formation in DTA^het^ mice, a 7-day dynamic histomorphometric analysis using calcein labeling was performed. The result showed that mineralized surface, mineral apposition rate (MAR) and bone formation rate (BFR) were significantly decreased in DTA^het^ mice (Figure 4E and F). Serum procollagen type 1 N-terminal propeptide (P1NP), a bone formation index, was also reduced after osteocyte ablation (Figure 4G). Meanwhile, *in vitro* osteogenesis showed there were less osteogenesis and mineralization in DTA^het^ mice compared to WT (Figure 4H and I), and the mRNA level of osteogenic markers including Alp, Ocn, Runx2 was also decreased (Figure 4J).

**Figure 4.**
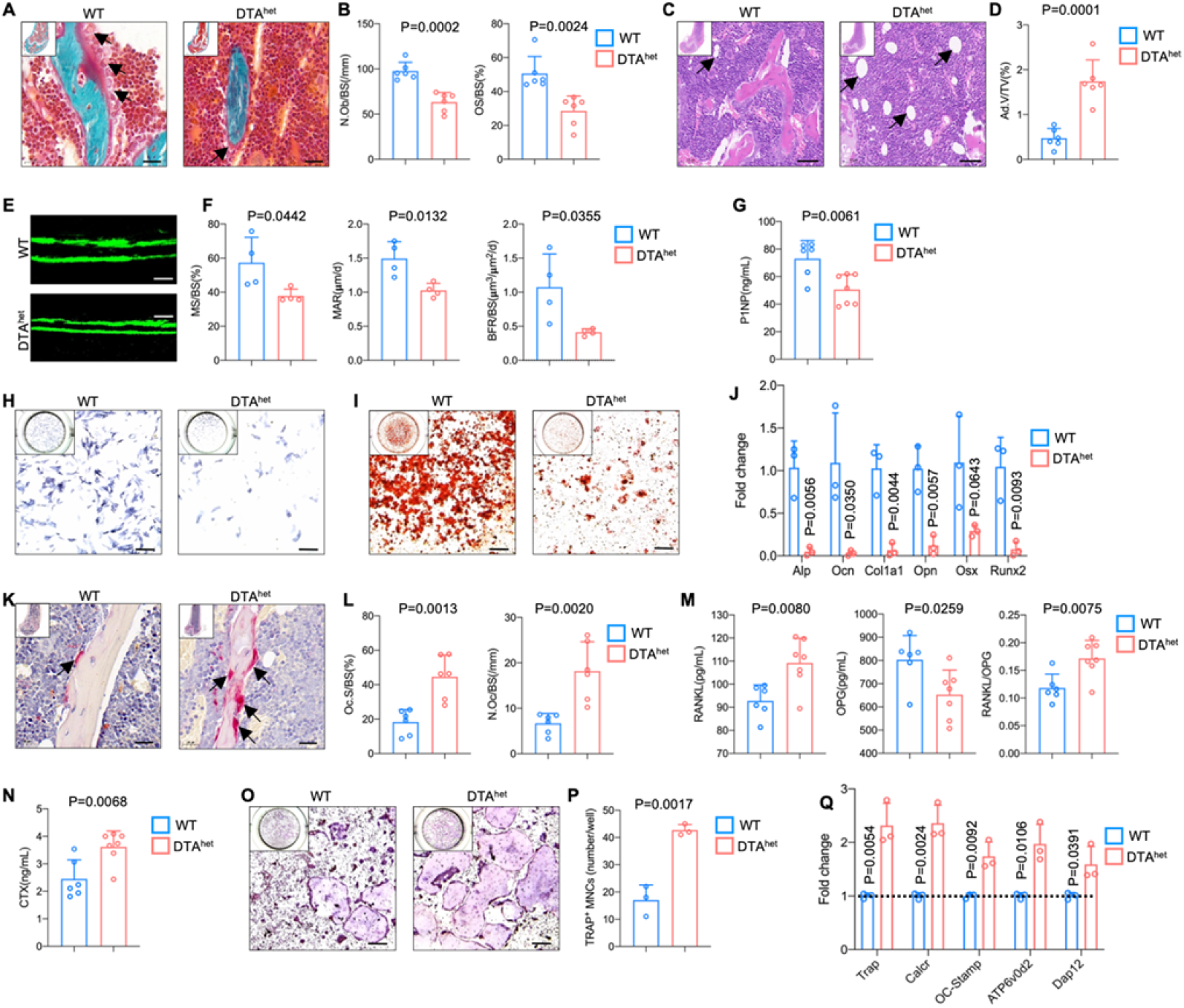
Ablation of osteocytes alters mesenchymal lineage commitment and promoted osteoclastogensis. (**A-B**) Goldner trichrome staining of male WT and DTA^het^ mice femur at 4 weeks (**A**) and histomorphometry analysis of osteoblast numbers (N.Ob/BS) (arrows) and osteoid-covered surface (OS/BS) (**B**) (n=6 per group). Scale bar, 20 μm. (**C-D**) Hematoxylin-eosin staining of WT and DTA^het^ mice femur at 4 weeks (**C**) and histomorphometry analysis of adipocyte (arrows) volume (Ad.V/TV) (**D**) (n=6 per group). Scale bar, 50 μm. (**E-F**) Representative images of calcein double labeling of the mineral layers of male WT and DTA^het^ mice femur at 4 weeks (**E**) and histomorphometry analysis of the mineral surface (MS/BS), mineral apposition rate (MAR) and bone formation rate (BFR/BS) (**F**) (n=4 per group). Scale bar, 50 μm. (**G**) ELISA of the concentration of bone formation index P1NP in the serum (n=6-7 per group). (**H-I**) Alp staining (**H**) and alizarin red staining (**I**) after osteoblast differentiation for 7 days and 21 days. Data are representative of three independent experiments. Scale bar, 250 μm. (**J**) RT-qPCR analysis of osteoblast signature genes expression at the mRNA level after osteoblast differentiation for 7 days (n=3 per group from three independent experiments), indicating impaired osteogenesis and increased adipogenesis in DTA^het^ mice. (**K-L**) TRAP staining of WT and DTA^het^ mice femur at 4 weeks (**K**) and histomorphometry analysis of osteoclast (arrows) surface (Oc.S/BS) and osteoclast numbers (N.Oc/BS) (**L**) (n=6 per group). Scale bar, 20 μm. (**M**) ELISAs of the concentration of RANKL, OPG and the ratio of RANKL/OPG in the serum (n=6-7 per group). (**N**) ELISA of the concentration of bone resorption index CTX in the serum (n=6-7 per group). (**O-P**) TRAP staining of after osteoclast differentiation for 5 days (**O**) and quantitative analysis (**P**) of TRAP positive cells (nucleus > 3) per well (n=3 per group from three independent experiments). Scale bar, 250 μm. (**Q**) RT-qPCR analysis of osteoclast signature genes expression at the mRNA level after osteoblast differentiation for 5 days (n=3 per group from three independent experiments), showing increased osteoclastogensis in DTA^het^ mice. Error bar represents the standard deviation.

In the aspect of osteoclastogenesis, histomorphometric analysis revealed that osteoclasts numbers and surface were significantly increased after osteocytes deletion (Figure 4K and L). Circulatory RANKL was also increased in DTA^het^ mice (Figure 4M). In contrast, circulatory osteoprotegrin (OPG), a decoy receptor of RANKL, was decreased, leading to the elevated ratio of RANKL/OPG (Figure 4M). Serum collagen type I c-telopeptide (CTX), a bone resorption index, was also significantly augmented in DTA^het^ mice compared to WT mice (Figure 4N), which implicated a high level of osteoclast activity of DTA^het^ mice in vivo. To assess the effects on osteoclasts after osteocyte ablation, bone marrow derived macrophages (Bmms) were collected respectively from DTA^het^ and WT mice and osteoclast differentiation was induced in vitro. The number of osteoclasts was substantially increased in DTA^het^ mice (Figure 4O and P). Also, the expression of the signature genes of osteoclasts including TRACP, Calcr, OC-stamp at the mRNA level was significantly upregulated in DTA^het^ mice (Figure 4Q). Together, osteocytes ablation impaired osteogenesis and promoted osteoclastogenesis.

### Alteration of hematopoietic lineage commitment by osteocyte ablation

As a part of the skeletal system, bone marrow has its vital functions in maintaining bone homeostasis(Divieti Pajevic and Krause 2019; Fulzele, et al. 2013; Asada, et al. 2013). HSCs give rise to lymphoid and myeloid lineage cells to establish the hematopoietic and immune system. To gain a full insight into the role of osteocyte in bone marrow homeostasis, single cell RNA sequencing (scRNA-seq) was performed using 10× Genomics Chromium platform. After rigorous quality control, gene expression data from 26562 cells (13835 and 12727 cells from 4-week littermate WT and DTA^het^ mice respectively) were compiled for clustering analysis, and there revealed 10 distinct populations visualized with uniform manifold approximation and projection (UMAP) embeddings (Figure 5A, B and C). These 10 distinct populations included B cell, hematopoietic stem cell and progenitor cell (HSPC), megakaryocyte, neutrophil, erythrocyte, monocyte, dendritic cell (DC), macrophage, T cell and mesenchymal stem cell (MSC) (Figure 5A and C). Proportion analysis revealed a significant expansion of neutrophils in DTA^het^ mice (Figure 5D and E). Also, the number of B cells was significantly less in DTA^het^ mice than that in WT mice (Figure 5D and E), which implicated that osteocytes ablation induced lymphoid-myeloid malfunction in the bone marrow. To further dissect the differences in the bone marrow development between two groups, RNA velocity was performed. The result showed that DTA^het^ mice have stronger directionality of velocity vectors from the HSPC population to the neutrophil population compared to WT mice (Figure 5F), implying that osteocytes deletion altered HSPC differentiation. Meanwhile, myeloid trajectory analysis revealed that there was a significantly higher pseudotime density distribution in G4 cell (a subcluster of neutrophil) in DTA^het^ mice (Figure 5G). In contrast, lymphoid trajectory analysis demonstrated a relatively lower pseudotime density distribution in pre-B cell and immature B cell (subclusters of B cell) in DTA^het^ mice (Figure 5H).

**Figure 5.**
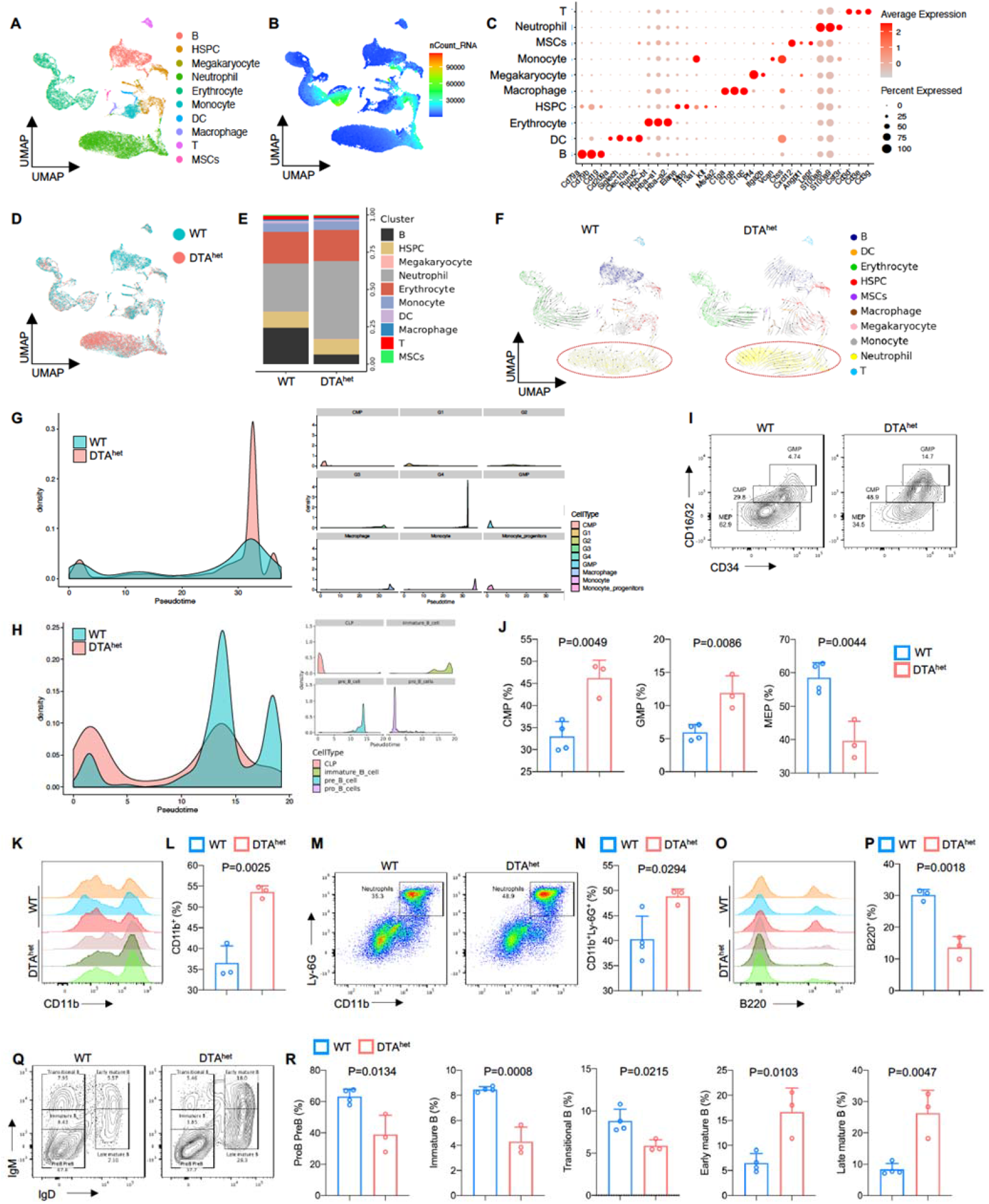
Alteration of hematopoietic lineage commitment by osteocyte ablation. (**A-B**) The UMAP plot of cells isolated from the bone marrow of 4 weeks WT and DTA^het^ mice and inferred cluster identity (**A**) and number of mRNA per cell (**B**). (**C**) Dot plot showing the scaled expression of selected signature genes for each cluster. Dot size represents the percentage of cells in each cluster with more than one read of the corresponding gene and dots are colored by the average expression of each gene in each cluster. (D-E) The UMAP plot of cells shown by sample (**D**) and proportions of each cluster in two samples (**E**). (F) RNA velocity analysis of clusters of WT and DTA^het^ mice shown by the UMAP embedding, showing stronger directionality of velocity vectors from HSPC cluster to neutrophil cluster in DTA^het^ mice. (**G**) Trajectory analysis of myeloid clusters of WT and DTA^het^ mice, demonstrating myeloid-biased hematopoiesis in DTA^het^ mice. (**H**) Trajectory analysis of lymphoid clusters of WT and DTA^het^ mice, demonstrating impaired lymphopoiesis in DTA^het^ mice. (**I-J**) Representative image of flow cytometry (**I**) and analysis of proportions of myeloid progenitors (CMP, GMP and MEP) (**J**) of 4 weeks WT and DTA^het^ mice (n=3-4 per group). (**K-L**) Representative image of flow cytometry (**K**) and analysis of proportions of CD11b^+^ myeloid cells (**L**) of 4 weeks WT and DTA^het^ mice (n=3 per group). (**M-N**) Representative image of flow cytometry (M) and analysis of proportions of neutrophils (**N**) of 4 weeks WT and DTA^het^ mice (n=3-4 per group). (**O-P**) Representative image of flow cytometry (**O**) and analysis of proportions of B220^+^ lymphoid cells (**P**) of 4 weeks WT and DTA^het^ mice (n=3 per group). (**Q-R**) Representative image of flow cytometry (**Q**) and analysis of proportions of ProB PreB, immature B, transitional B, early mature B and late mature B (**R**) of 4 weeks WT and DTA^het^ mice (n=3-4 per group), indicating altered B cell development pattern in DTA^het^ mice. Error bar represents the standard deviation.

To corroborate the results observed from scRNA-seq, flow cytometry and further analysis were performed after removing adherent cells (Figure 5 - figure supplement 1A and B). Although there was no significant change of hematopoietic stem cell (HSC) (Lin^-^c-Kit^+^Sca1^+^, LSK^+^ cell) numbers between DTA^het^ and WT mice (Figure 5 - figure supplement 2A and B), DTA^het^ mice demonstrated significantly increased number of short-term HSC (ST-HSC) with decreased number of long-term HSC (LT-HSC), indicating that HSC in DTA^het^ mice bone marrow was mobilized (Figure 5 - figure supplement 2C and D). Further flow cytometry analysis revealed that the number of myeloid progenitors including common myeloid progenitors (CMP), granulocyte-monocyte progenitors (GMP) and common monocyte progenitors (cMoP) were substantially increased after osteocyte ablation (Figure 5I and J, Figure 5 - figure supplement 2E and F), and megakaryocyte erythroid progenitors (MEP) numbers were decreased (Figure 5I and J). Meanwhile, total CD11b^+^ myeloid cells were also increased (Figure 5K and L) in DTA^het^ mice, in which both neutrophil and monocytes significantly expanded (Figure 5M and N, Figure 5 - figure supplement 2G and H). In addition, while the proportion of common lymphoid progenitors (CLP) was not altered in DTA^het^ mice (Figure 5I and J), total B220^+^ lymphoid cells reduced remarkably after osteocyte ablation (Figure 5O and P), in which DTA^het^ mice showed a relatively lower proportion of early B cell (pro-B pre-B, immature B and transitional B cell) and a relatively higher proportion of late B cell (early mature B and late mature B) (Figure 5K and L), which suggested that B cell development was impaired along the immature B to mature B cell transition in DTA^het^ mice. As scRNA-seq revealed that neutrophil underwent a significant change after osteocyte ablation, neutrophil population were further reclustered into four subclusters from G1 to G4 (Figure 5 - figure supplement 3A and B) and G4 population was significantly increased in DTA^het^ mice compared to WT mice (Figure 5 - figure supplement 3C and D), which implied that osteocyte ablation accelerated neutrophil maturation. Consistent with this observation, neutrophil functions including activation, chemotaxis were all upregulated in DTA^het^ mice (Figure 5 - figure supplement 3E and F). Genes related to glycolysis and necroptosis were also upregulated (Figure 5 - figure supplement 3G and H), indicating that osteocyte ablation induced neutrophil functions. Together, these results demonstrated that osteocyte ablation altered hematopoietic lineage, characterized by the shift from lymphopoiesis to myelopoiesis.

### Organismal senescence of osteoprogenitors and myeloid lineage cells leads to the skeletal premature aging

Senescence occurred during development as a precise programmed cellular process, contributes to cell fate specification, tissue patterning and transient structure removal(Munoz-Espin and Serrano 2014; Rhinn, Ritschka, and Keyes 2019). Given that DTA^het^ mice had a skeletal premature aging with increased myelopoiesis, osteoporosis and sarcopenia, we hypothesized that osteocyte ablation may induce organismal senescence of osteoprogenitors and myeloid lineage cells. ScRNA-seq revealed that total bone marrow had increased senescence with a higher senescence associated secretory phenotype (SASP) score in DTA^het^ mice compared to WT mice (Figure 6A). DTA^het^ mice also had increased maturity in bone marrow reflected from RNA velocity (Figure 6B). Meanwhile, circulatory SASP including Tnf-*α*, Il1β and Il6 were also elevated in DTA^het^ mice (Figure 6C). Further scRNA-seq analysis uncovered that mesenchymal stem cell (MSC), CMP, monocyte and its subcluster Ly6c2 monocyte, neutrophil and its subcluster G2, G3 and G4 had increased SASP scores (Figure 6D-G). RT-qPCR also verified the elevated senescence with increased gene expressions including p16 and p21 in DTA^het^ mice (Figure 6 - figure supplement 1A and B). Together, these results suggested that osteocyte reduction induced senescence in osteoprogenitors and myeloid lineage cells.

**Figure 6.**
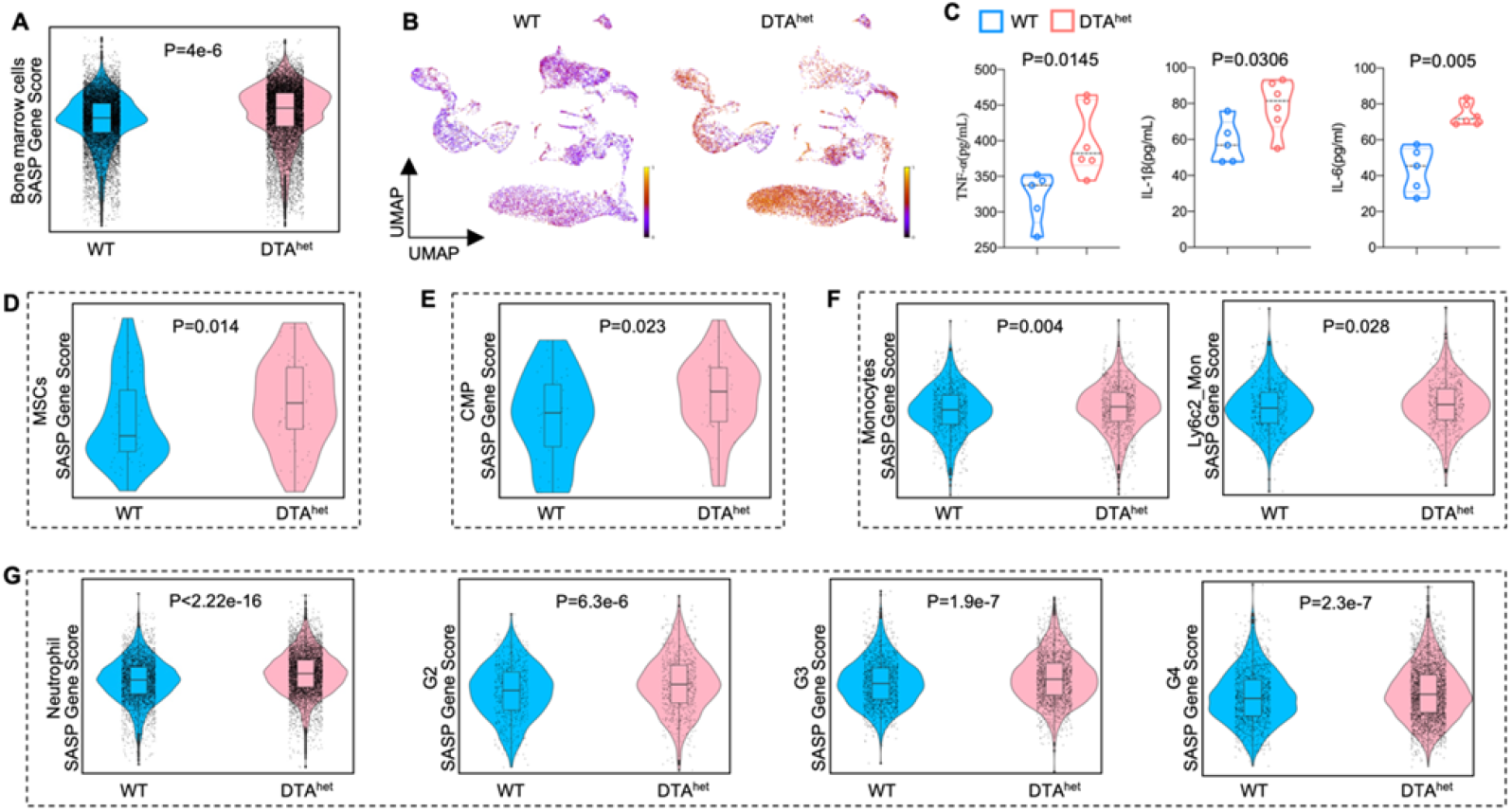
Organismal senescence of osteoprogenitors and myeloid lineage cells leads to the skeletal premature aging. (**A**) Comparisons of total bone marrow cells SASP score between 4 weeks WT and DTA^het^ mice. (**B**) Latent time of RNA velocity analysis of WT and DTA^het^ mice shown by the UMAP embedding. (**C**) ELISAs of the concentration of TNF-*α*, IL-1*β* and IL-6 of 4 weeks WT and DTA^het^ mice in the serum (n=5-6 per group). (**D**) Comparisons of MSCs SASP score between 4 weeks WT and DTA^het^ mice, indicating the senescence of osteoprogenitors in DTA^het^ mice. (**E**) Comparisons of CMP SASP score between 4 weeks WT and DTA^het^ mice. (**F**) Comparisons of monocytes and its subcluster Ly6c2^+^ monocytes SASP score between 4 weeks WT and DTA^het^ mice. (**G**) Comparisons of neutrophils and its subcluster (G2, G3 and G4) SASP score between 4 weeks WT and DTA^het^ mice, indicating the senescence of myeloid lineage cells. Error bar represents the standard deviation.

Owning to the fact that osteoblast derived from mesenchymal stem cell lineage, we next investigated whether accumulation of osteoprogenitor cell senescence impaired osteogenesis. GO analysis revealed that downregulated genes after osteocyte ablation were enriched in ossification and biomineral tissue development (Figure 6 - figure supplement 1C), which was consistent with the finding of impaired osteoblast differentiation (Figure 4H-J). Similarly, Kyoto Encyclopedia of Genes and Genomes (KEGG) analysis revealed that the subcluster 2 and 4 of Ly6c2^+^ monocytes demonstrated the enrichment of osteoclast differentiation related genes after osteocyte ablation (Figure 6 - figure supplement 1D and E), which was corroborated in our enhanced in vitro osteoclast differentiation (Figure 4O-Q). Together, our data suggested that senescence in osteoprogenitors and myeloid lineage cells led to the impaired osteogenesis and increased osteoclastogenesis, respectively.

## Discussion

In this study, we demonstrated an important role of osteocytes in regulating organismal senescence of bone and bone marrow. Partial ablation of osteocytes^DMP-1^ caused severe sarcopenia, osteoporosis and degenerative kyphosis, which led to shorter lifespan. Acquisition of a senescence-associated secretory phenotype (SASP) in both osteoprogenic and myeloid lineage cells is underlying cause that led to the skeletal premature aging phenotype of impaired osteogenesis, increased osteoclastogenesis and myelopoiesis.

Sarcopenia usually occurs concurrently with osteoporosis during aging(Clynes et al. 2021). Our study has showed for the first time that osteocyte ablation caused severe sarcopenia and muscle atrophy. In consistent with our observation, previous studies have reported that osteocyte-specific ablation of Cx43 impaired muscle formation(Shen et al. 2015). Osteocyte-derived factors has also been shown to stimulate myogenic differentiation in vitro(Huang et al. 2017). On the contrary, specific deletion of Mbtps1 in osteocyte promotes soleus muscle regeneration and increase its size with age(Gorski et al. 2016). Sclerostin, an osteocyte-derived circulating protein, is negatively correlated with skeletal muscle mass(Kim et al. 2019). Previously there has been a study showing weak DMP-1 expression in skeletal muscle fibers(Lim et al. 2017). This has led us to suggest that sarcopenia may be caused directly by the DMP-1 expression in muscle. However, our histology finding of no obvious changes in the total number of nuclei of muscle in partial ablation of DMP-1 positive osteocytes suggested that the sarcopenia and muscle atrophy phenotype is most likely caused by the disturbance of osteocyte-muscle crosstalk. Certainly, further studies based on a more specific osteocyte ablation model are needed to understand the link of osteocytes between osteoporosis and sarcopenia. Nevertheless, severe kyphosis observed in these osteocyte ablation mice, support our hypothesis of direct osteocyte-muscle crosstalk, as kyphosis is the direct result of the significant bone loss and sarcopenia(Wijshake et al. 2012; Woods et al. 2020).

Osteocytes regulate the process of bone resorption mediated by osteoclasts and coupled bone formation mediated by osteoblasts via secreting products like sclerostin and RANKL(Tresguerres, et al. 2020; van Bezooijen et al. 2005; Nakashima et al. 2011). Theoretically, osteocyte ablation may lead to lower expression of sclerostin and RNAKL with increased osteogenesis and impaired osteoclastogenesis. But our results demonstrated that osteocyte ablation impaired osteogenesis and induced osteoclastogenesis. Furthermore, the expression of sclerostin mRNA was reduced as expected, the serum RNAKL was increased after osteocyte ablation. We speculated that induction of SASP in both osteoprogenitors and myeloid progenitors may be account for the underlying cause. Senescent osteoprogenitors have reduced self-renewal capacity and predominantly differentiate into adipocytes as opposed to osteoblasts(Chen, et al. 2016; Li et al. 2017; Rosen et al. 2009). Consistently, our model indicated an increased adipogenesis after osteocyte ablation. Also, fat-induction factors inhibit osteogenesis during adipogenesis(Chen, et al. 2016). Thus, osteocyte ablation induced senescence accumulation in osteoprogenitors leading to the cell commitment towards adipogenesis with impaired osteogenesis. As for enhanced osteoclastogenesis, besides the production of RANKL from osteogenic cell like osteocytes and osteoblasts(Nakashima, et al. 2011; Fumoto et al. 2014), other cells like adipocyte, T cell also secret RANKL to regulate bone metabolism(Yu et al. 2021; Hu et al. 2021; Djaafar et al. 2010; Takayanagi et al. 2000). Also, B cell can produce OPG to regulate RANKL/OPG axis(Li et al. 2007). In our model, increased adipogenesis, T cell expansion (data not shown) and decreased B cell number may compensate for the altered RANKL/OPG axis.

Bone marrow, embedded in the skeletal system, has a close link with matrix-embedded osteocyte. Previous studies have reported that osteocyte regulates myelopoiesis via Gs*α* -dependent and -independent signaling(Fulzele, et al. 2013; Azab, et al. 2020). Recent study also reported that osteocyte mTORC1 signaling regulates granulopoiesis via secreted IL-19(Xiao, et al. 2021). Meanwhile, sclerostin secreted by osteocyte adversely affects B cell survival(Horowitz and Fretz 2012). In our study, when osteocytes were partially depleted, myelopoiesis especially granulopoiesis was significantly induced, but B cell development was significantly impaired. Further studies demonstrated that HSC was mobilized and shifted to myelopoiesis with increased CMP, GMP, cMoP and CD11b^+^ myeloid cells, in which monocytes and neutrophils were increased, and neutrophil function was also activated after osteocyte ablation. While B cell number was severely reduced with altered development pattern. Interestingly, previous study has shown that osteoblastic cell support megakaryopoiesis and platelet formation(Xiao et al. 2017). In our study, the number of MEP (erythrocyte and platelet precursors) was also reduced, and scRNA-seq analysis showed no significant change in erythrocyte population (data not shown), inferring that osteocyte may also participate in regulating platelet formation. Together, these results provided evidence that osteocyte play essential roles in maintaining the HSC niche homeostasis. In conclusion, we demonstrated a critical role of osteocytes in regulating organismal senescence of bone, and bone marrow (Figure 7). Ablation of osteocytes induced SASP accumulation in bone marrow osteoprogenitors and myeloid lineage cells, which altered MSC and HSC lineage commitments with impaired osteogenesis, promoted myelopoiesis and osteoclastogenesis, leading to the skeletal premature aging phenotype with severe sarcopenia, osteoporosis, degenerative kyphosis and bone marrow myelopoiesis, thus shortened lifespan of mice. Targeting osteocyte function and cell fate may shed light on the therapeutic regimens for aging associated bone diseases.

**Figure 7.**
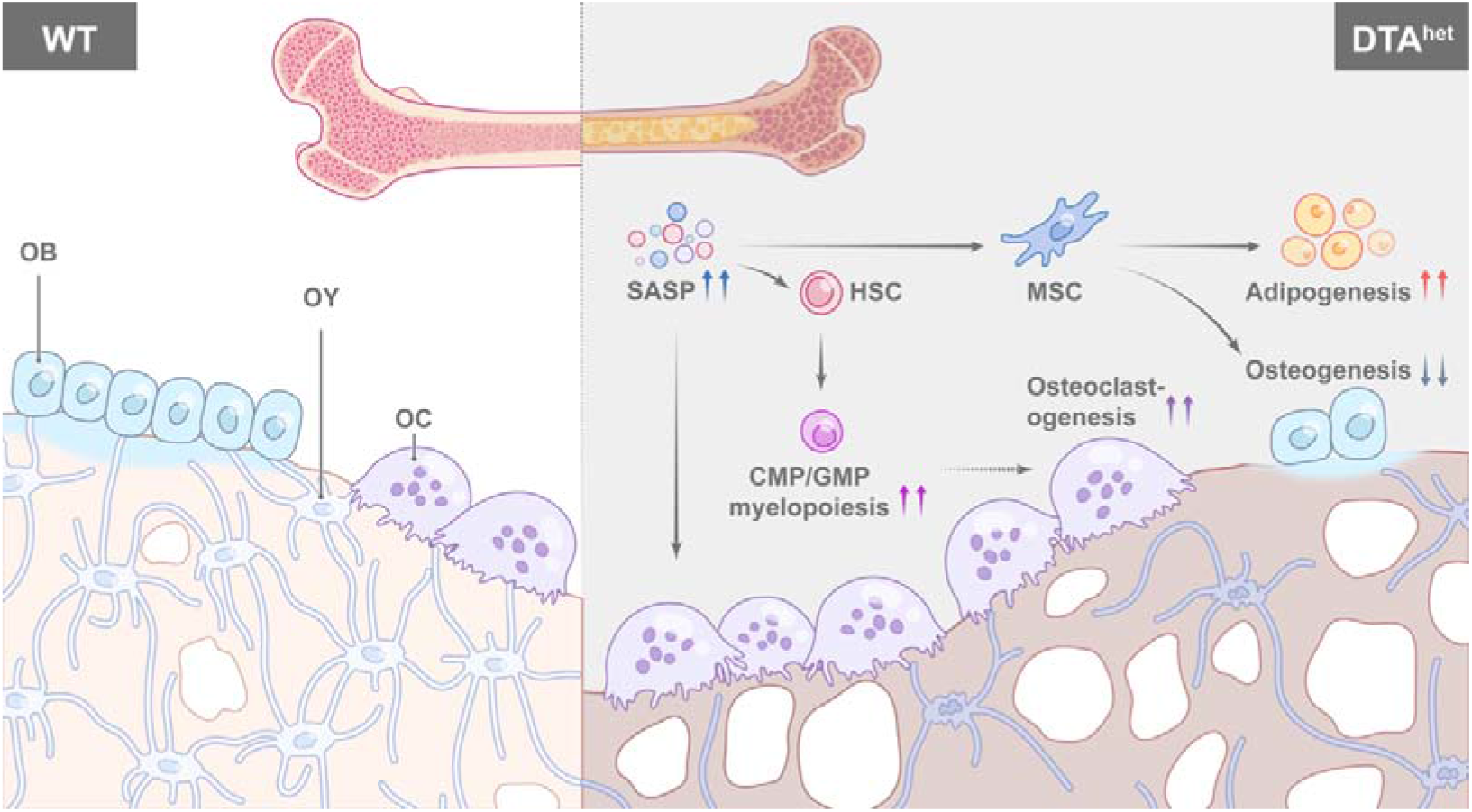
Schematic diagram of osteocyte ablation induced skeletal senescence. Ablation of osteocytes induced SASP accumulation in bone marrow osteoprogenitors and myeloid lineage cells, which altered MSC and HSC lineage commitments with promoted adipogenesis, myelopoiesis and osteoclastogenesis at the expense of osteogenesis and lymphopoiesis, leading to the skeletal premature aging phenotype with severe sarcopenia, osteoporosis, degenerative kyphosis and bone marrow myelopoiesis, thus shortened lifespan of mice.

## Materials and methods

### Mice

All mouse lines were maintained on a C57BL/6J background. DMP-1^cre^ mice were provided by J. Q. (Jerry) Feng from Texas A&M College of Dentistry, USA (Jackson Laboratory stock number, 023047). DTA^fl/+^ mice were from GemPharmatech (strain ID, T009408). Osteocyte ablation mice model during development was established by crossing DMP-1^cre^ mice with DTA^fl/fl^ mice to obtain DMP-1^cre^ DTA^fl/+^ mice (DTA^het^). All mice experiments were approved by the Animal Care and Use Committee of Shanghai Jiao Tong University Affiliated Sixth People’s Hospital.

### Bone histomorphometry analysis

Mice femur was dissected and fixed in 4% paraformaldehyde (PFA) for two days and further decalcified with 10% EDTA (pH=7.2) in 4°C for about 2 weeks. Then specimens were embedded in paraffin and sectioned at 4 μm thickness. TRAP staining was performed for osteoclast analysis. H&E staining was performed for adipocyte and osteocyte analysis. For osteoblast analysis, undecalcified femur was embedded in plastic and sectioned at 5 μm thickness and Goldner trichrome staining was performed. For dynamic histomorphometry analysis, double calcein-labeling was used. Briefly, each mouse was given 30 μg/gram body weight Calcein (Sigma) on day 1 and day 7 by intraperitoneal injection before sacrifice. Bones were then fixed, dehydrated, embedded in plastic and cut into 5μm slices and calculated using the software under fluorescence. Bioquant Osteo software (Bioquant) was used for histomorphometry analysis. Accepted nomenclature was used to report the results(Dempster et al. 2013). ImageJ was used to measure the number of osteocyte lacunae.

### Immunofluorescence staining

Both ends of the mice tibias/femurs were removed. Then they were embedded in OCT for frozen sectioning and cut parallel to the long axis of the long bones. Stop cutting when the maximum cross section of the long bones was observed. The OCT around the rest of the bones were melted at room temperature. The bone samples remained were washed 3 times in PBS for 10 minutes and fixed in 4% paraformaldehyde (PFA) for 2 hours. Then, they were immersed in 0.1%Triton X-100 for 1 hour, blocked using 3% BSA and stained using Alexa Fluor™ 568 Phalloidin (Invitrogen) for 48 hours at 4°C in the dark with gentle shake. The samples were washed 3 times with PBS for 10 minutes. The cross section of the sample was inverted in the confocal dish. Pictures were captured using confocal microscopy (Olympus) and ImageJ was used to measure the number dendrites per osteocyte.

### Bone density measurements

Mice femurs and L3 lumbar were stripped of soft tissue and fixed in 4% PFA overnight at 4 °C, then stored in 70% ethanol until scanned using the μCT instrument (SkyScan 1176). Relevant structure parameters of the μCT instrument were as previous reported(Ding et al. 2022): scanning voxel size, 9×9×9 um^3^; X-ray tube potential, 50 kV and 450 uA; integration time, 520 ms; rotation Step, 0.4° for 180° scanning. CTAn micro-CT software version 1.13 (Bruker) was used to analyze the images. The threshold value (grayscale index) for all trabecular bone was 75. For all cortical bone the threshold value (grayscale index) was 110. The femurs were analyzed at a resolution of 9 μm. The volumetric regions for trabecular analyses include the secondary spongiosa located 1 mm from the growth plate and extending 1.8 mm (200 sections) proximally. For cortical bone analysis, the volumetric regions include 600 μm long at mid-diaphysis of the femur (300 μm extending proximally and distally from the diaphyseal midpoint between the proximal and distal growth plates). For vertebrae, the volumetric regions include the entire trabecular region without the primary spongiosa (300μm below the cranial and above the caudal growth plate). Morphometric parameters including bone mineral density (BMD), bone volume/total volume fraction (BV/TV), trabecular number (Tb.N), trabecular thickness (Tb.Th), trabecular separation (Tb.Sp), cortical thickness(Ct.Th) and cortical porosity(Ct.Po) were calculated.

### Gait analysis

CatWalk automated gait analysis system (Noldus Information Technology) was used to analysis gait. Mice were expected to run along a special glass plate with a green LED lit and a high-speed video camera under it. Their paws were captured by the camera. Before the formal experiments, the mice were habituated in the plate to achieve an unforced locomotion. Three compliant runs without stopping, changing direction and turning around were analyzed with Catwalk Software. Relevant data were generated by Catwalk Software after each footprint was checked manually. Data including stride length, swing speed and normal step sequence radio were analyzed.

### Whole mount alcian blue/alizarin red staining

The skin and viscera of the intact fetal mice (E19.0) were removed. The embryos were fixed in 95% ethanol overnight and then degreased in absolute acetone overnight with gentle agitation. The embryos were stained overnight in 0.015% alcian blue (Sigma) /0.005% alizarin red (Sigma) in 70% ethanol with gentle agitation. They were washed in 70% ethanol for 30 min three times and digested using 1% KOH solution. When most of the soft tissue was digested, the embryos were immersed in 75% (vol/vol)1% KOH/glycerol solution for further clearing. Graded glycerol was changed according to the degree of embryos digestion and relevant pictures were obtained under the microscope (Leica).

### Whole-body μCT scan

13- and 37-week-old DTA^het^and wild-type mice were deeply anesthetized and carefully positioned with a dedicated cradle and holder to capture the whole-body (excluding the tail) radiographs at a resolution of 35 μm using the μCT instrument (SkyScan 1176). Scanning details were listed as following: X-ray tube potential, 65 kV and 375 uA; exposure time, 150 ms; rotation step, 0.5° for 180° scanning. CTAn micro-CT software version 1.13 (Bruker) was used to reconstruct pictures.

### RNA-seq

Total RNA of whole bone with bone marrow flushed out from 4 weeks WT and DTA^het^ mice was extracted using Trizol reagent (Thermofisher), quantified and purified using Bioanalyzer 2100 and RNA 6000 Nano LabChip Kit (Agilent). Following purification, mRNA library was constructed, fragmented, amplified and loaded into the nanoarray and sequencing was performed on Illumina Novaseq™ 6000 platform following the vendor’s recommender protocol. After sequencing, generated reads were filtered and mapped to the reference genome using HISAT2 (v2.0.4) and assembled using StringTie (v1.3.4d) with default parameters. Then, all transcriptomes from all samples were merged to reconstruct a comprehensive transcriptome using gffcompare software (v0.9.8) and the expression levels of all transcripts were calculated by Stringtie and ballgown. Differential gene analysis was performed by DESeq2 software and then subjected to enrichment analysis of GO functions. The data were deposited into the GEO repository (GSE202356, secure token for reviewer: ipqryuycnloznsz)

### Cell culture

#### In vitro osteoclastogenesis assay

The bone marrow of mice femurs and tibias were flushed to get bone marrow cells. Cells were cultured overnight by using α-MEM (Hyclone) which contains 10% FBS (Gibco), 100 μg/ml streptomycin (Gibco) and 100 U/ml penicillin (Gibco). The non-adherent cells were collected, layered on Ficoll-Paque (GE Healthcare) and separated through density gradient centrifugation at 4 °C and 2000 rpm for 20 min. The bmms were in the middle layer of the separation. Bmms were collected and washed twice with ice-cold PBS. To induce osteoclast differentiation, bmms (2.5*10^4^ cells per well for 96-well plates and 8*10^5^ per well for 6-well plates) were cultured by using α-MEM which contains 10% FBS, 100 μg/ml streptomycin, 100 U/ml penicillin, 100 ng/ml M-CSF (Peprotech) and 100 ng/ml RANKL (Peprotech) for 5 days before TRAP staining. Cells were cultured at 37°C in a humidified incubator at 5% CO_2_. The medium was changed every 2 days. At the end of assay (the fifth day), the cells were fixed and stained with Tartrate-resistant acid phosphatase (TRAP) kit according to the manufacturer’s instructions (Sigma) to quantify osteoclast numbers, or RNA was extracted as recommended protocol. TRAP-positive cells which contains more than three nuclei were counted as mature osteoclast-like cells (OCLs). The assay was repeated three times and number of OCLs per well were recorded for each biological replicate.

#### Harvest of calvarial osteoblasts and osteogenic differentiation

Neonatal DTA^het^ and WT pubs (P0) was euthanized and decapitated using scissors. The calvaria were separated and any loose connective tissue from the calvaria were removed. Then, the calvaria were digested five times using α-MEM which contains 0.1% collagenase (Roche) and 0.2% dispase (Roche) in a 37 °C constant temperature shaking table set at 250 rpm for 10 min. The last four times’ digestive production which contains calvarial osteoblasts were collected and cultured in α-MEM containing 10% FBS, 100 U/ml penicillin, 100 μg/ml streptomycin. After culturing the cells to 70–80 % confluence prior, they were re-plated at a density of 5000 cells per well for 96-well plates or 2×10^5^ cells per well for 6-well plates. When the cells were cultured to 70-80 % confluence prior, the medium was replaced with osteogenic differentiation medium (Cyagen) and changed every 2 days. After a week of differentiation, the cells were either fixed and ALP staining was performed, or RNA extraction was performed. After three weeks of differentiation, alizarin red staining was performed.

### RT-qPCR

Total RNA was isolated using RNeasy^®^ Mini Kit (Qiagen). 500 ng of total RNA was reverse transcribed into cDNA using PrimeScript™ RT Master Mix (Takara, RR036A). qPCR analyses were performed using SYBR Premix Ex Taq™ II (Takara, RR820L) and samples were run on the ABI HT7900 platform (Applied Biosystems). SYBR Green PCR conditions were 1 cycle of 95°C for 30 seconds, and 40 cycles of 95°C for 5 seconds and 34°C for 60 seconds. Melting curve stage was added to check primers specificity. Relative gene expression levels were calculated using the threshold cycle (2^-ΔΔCT^) method. Relevant primers were listed as below: Gapdh: 5’-ACC CAG AAG ACT GTG GAT GG-3’and 5’-CAC ATT GGG GGT AGG AAC AC-3’; p21: 5’-GTC AGG CTG GTC TGC CTC CG-3’ and 5’-CGG TCC CGT GGA CAG TGA GCA G-3’; p16: 5’-GTC AGG CTG GTC TGC CTC CG-3’ and 5’-CGG TCC CGT GGA CAG TGA GCA G-3’; Il6: 5’-CTG GGA AAT CGT GGA AT-3’ and 5’-CCA GTT TGG TAG CAT CCA TC-3’; Mcp1: 5’-GCA TCC ACG TGT TGG CTC A3’ and 5’-CTC CAG CCT ACT CAT TGG GAT CA-3’; Tnf: 5’-ATG AGA AGT TCC CAA ATG GC-3’ and 5’-CTC CAC TTG GTG GTT TGC TA-3’; Il1b: 5’-GCC CAT CCT CTG TGA CTC AT-3’ and 5’-AGG CCA CAG GTA TTT TGT CG-3’; Alp: 5’-TCA GGG CAA TGA GGT CAC AT-3’ and 5’-CCT CTG GTG GCA TCT CGT TA-3’; Ocn: 5’-CCC TGA GTC TGA CAA AGC CT-3’ and 5’-GCG GTC TTC AAG CCA TAC TG-3’; Col1a1: 5’-ATA AGT CCC TTC CTG CCC AC-3’ and 5’-TGG GAC ATT TCA GCA TTG CC-3’; Opn: 5’-ATG CCA CAG ATG AGG ACC TC-3’ and 5’-CCT GGC TCT CTT TGG AAT GC-3’; Osx: 5’-TCG GGG AAG AAG AAG CCA AT-3’ and 5’-CAA TAG GAG AGA GCG AGG GG-3’; Runx2: 5’-GCC CAG GCG TAT TTC AGA TG-3’ and 5’-GGT AAA GGT GGC TGG GTA GT-3’; Dmp1: 5’-CAG TGA GGA TGA GGC AGA CA-3’ and 5’-CGA TCG CTC CTG GTA CTC TC-3’; Sost: 5’-GCC GGA CCT ATA CAG GAC AA-3’ and 5’-CAC GTA GCC CAA CAT CAC AC-3’; Trap: 5’-TGG ACA TGA CCA CAA CCT GCA GTA-3’and 5’-TCG CAC AGA GGG ATC CAT GAA GTT-3’; Calcr: 5’-AGC CAC AGC CTA TCA GCA CT-3’and 5’-GAC CCA CAA GAG CCA GGT AA-3’; OC-Stamp: 5’-TGG GCC TCC ATA TGA CCT CGA GTA G-3’and 5’-TCA AAG GCT TGT AAA TTG GAG GAG T-3’; ATP6v0d2: 5’-ACA TGT CCA CTG GAA GCC CAG TAA-3’and 5’-ATG AAC GTA TGA GGC CAG TGA GCA-3’; Dap12:5’-CTG GTG TAC TGG CTG GGA TT-3’and :5’-CTG GTC TCT GAC CCT GAA GC-3’. All these primers were synthesized by Sangon Biotech company (Shanghai).

### Flow cytometry

Bone marrow cells were isolated by flushing the bone marrow of mice femurs and tibias with PBS and were dissociated into a single cell suspension by gently filtering them through 70 μm nylon mesh. After red blood cells lysis, the isolated cells were blocked by anti-mouse CD16/32 antibody (Biolegend, 101302) for 15 min and stained with fluorescence-conjugated antibodies for 30 min at 4°C in the dark. Relevant antibodies were listed as below and their catalog numbers were provided in the brackets:anti-Ly-6C-Pacific Blue™ (128013), anti-Ly-6C-PE (128007), anti-Ly-6G-Pacifìc Blue™ (127611), anti-Ly-6G-PE/Cy7 (127617), anti-CD16/32-FITC (101305), anti-CD115-PE (135505), anti-CD117-PE (105808), anti-CD117-APC/Cy7 (105825), anti-CD45R-PE/Cy5 (103209), anti-CD45R-APC (103212), anti-Ly-6A/E-APC (108111), anti-Ly-6A/E-Alexa Fluor^®^700(108142), anti-CD34-PerCP/Cyannine5.5 (128607), anti-CD135-APC (135309), anti-lineage cocktail-Pacifìc Blue™ (133305), anti-CD127-PE (121111), anti -CD127-APC(135011), anti-CD11b-FITC (101205) and anti-CD24-Pacifìc Blue™ (101819). All these antibodies were purchased from Biolegend. Samples were analyzed using cytometer CytoFlex (Beckman Coulter) and FlowJo software version 10.4. 50000 events were collected for each sample.

### Preparation of mice serum

For serum collection, mice were anesthetized with isoflurane and blood samples were collected from the ophthalmic vein. Samples were then centrifuged at 5000 rpm for 5 min. Supernatants were transferred to a new tube and centrifuged at 5000 rpm for 5 min again. Supernatants were collected to a new tube and treated with liquid nitrogen fast and then stored at −80 °C.

### Enzyme-linked immunosorbent assay (ELISA)

Elisa was performed as kit instructions (Jianglai). Briefly, working standards and diluted samples were prepared and added to each well. Plates were sealed and incubated for 1 hour at 37 °C. After washing three times, 100 μl enzyme-labeled reagents were added and plates were incubated for 1 hour at 37 °C. Finally, TMB substrates were added and incubated for 15-30 minutes at 37 °C followed by Stop solution addition. Then plates were read at 450 nm within 5 minutes.

### Singe cell collection, library construction and sequencing

Bone marrow cells from WT and DTA^het^ mice were flushed and sieved through a 70 μm cell strainer. After red blood cell analysis, dissociated single cells were stained with AO/PI for viability assessment. Single-cell RNA sequencing (scRNA-seq) was performed using 10× Genomics Chromium platform. Related operations including Generation of gel beads in emulsion (GEMs), barcoding, GEM-RT cleanup, complementary DNA amplification and library construction were all carried out following the manufacturer’s protocol. By using 150-base-pair paired-end reads, the final libraries were sequenced on the Illumina NovaSeq 6000 platform. The scRNA-seq data could be accessed from GEO database (GSE202516, secure token for reviewer: ihudckqqxvopruz)

### Data processing, dimension reduction, unsupervised clustering and annotation

ScRNA-seq data analysis was performed by NovelBio Co.,Ltd with NovelBrain Cloud Analysis Platform (www.novelbrain.com). Fastp was applied with default parameters filtering the adaptor sequence and the low-quality reads were removed to achieve the clean data. Then the feature-barcode matrices were obtained by aligning reads to the mouse genome (mm10 Ensemble: version 92) using CellRanger v3.1.0. Down sample analysis among samples sequenced was applied according to the mapped barcoded reads per cell of each sample and finally achieved the aggregated matrix. Cells contained over 200 expressed genes and mitochondria UMI rate below 20% passed the cell quality filtering and mitochondria genes were removed in the expression table.

Seurat package (version: 3.1.4, https://satijalab.org/seurat/) was used for cell normalization and regression based on the expression table according to the UMI counts of each sample and percent of mitochondria rate to obtain the scaled data. PCA was constructed based on the scaled data with top 2000 high variable genes and top 10 principals were used for tSNE construction and UMAP construction. Utilizing graph-based cluster method, the unsupervised cell cluster results based the PCA top 10 principal were acquired, and the marker genes by FindAllMarkers function with wilcox rank sum test algorithm was calculated under following criteria: lnFC > 0.25, p value < 0.05 and min.pct > 0.1. To identify the cell type detailed, the clusters of same cell type were selected for re-tSNE analysis, graph-based clustering and marker analysis.

### Identification of differential gene expression and gene enrichment analysis

To identify differentially expressed genes among samples, the function FindMarkers with wilcox rank sum test algorithm was used under following criteria: lnFC > 0.25, p value < 0.05 and min.pct > 0.1. Gene ontology (GO) analysis was performed to facilitate elucidating the biological implications of marker genes and differentially expressed genes. The GO annotations from NCBI (http://www.ncbi.nlm.nih.gov/), UniProt (http://www.uniprot.org/) and the Gene Ontology (http://www.geneontology.org/) were downloaded. Fisher’s exact test was applied to identify the significant GO categories and FDR was used to correct the p-values. Pathway analysis was used to find out the significant pathway of the marker genes and differentially expressed genes according to KEGG database. Fisher’s exact test was applied to select the significant pathway, and the threshold of significance was defined by P-value and FDR. To characterize the relative activation of a given gene set such as pathway activation, QuSAGE (2.16.1) analysis was performed, and related gene set involving neutrophil function and SASP were from ref. (Xie et al. 2020; Zhang et al. 2021) and listed in supplement file table 2.

### Developmental trajectory inference and RNA velocity analysis

The Single-Cell Trajectories analysis was applied utilizing Monocle2 (http://cole-trapnell-lab.github.io/monocle-release) using DDR-Tree and default parameter. Before Monocle analysis, marker genes of the Seurat clustering result and raw expression counts of the cell passed filtering were selected. Based on the pseudo-time analysis, branch expression analysis modeling (BEAM Analysis) was applied for branch fate determined gene analysis. To estimate the cell dynamics, RNA Velocity analysis was performed through scVelo package (Version0.2.3) based on ScanPy package (Versionv1.5.0) with default parameters.

### Statistical analysis

All data were analyzed using GraphPad Prism (v8.2.1) software for statistical significance. P value was determined by the student’s t test for two-group or one-way ANOVA test for multiple group comparisons. Gehan-Breslow-Wilcoxon test was used for analyzing Kaplan-Meier curve of WT and DTA^het^ mice.

## Author contributions

J.J.G., C.Q.Z. and. M.H.Z. conceived, designed, and supervised the study. P. D., C.A.G., Y.S.G. performed the experiment and analyzed the data. D.L.L., H.L., J.X., X.Y.C., Y.G.H provided suggestions. P. D., C.A.G. wrote the manuscript.

## Acknowledgements

This work was supported by National Natural Science Foundation of China (82002339 to J.J.G, 81820108020 to C.Q.Z.) and Shanghai Frontiers Science Center of Degeneration and Regeneration in Skeletal System (BJ1-9000-22-4002).

## Data and materials availability

ScRNA-Seq and RNA-seq data have been deposited into GEO repository with accession codes GSE202516 and GSE202356 respectively. Additional data that support the findings of this study are available from the corresponding author on request. Source data are provided with this paper.

## Supplementary figure legend

**Figure 2 - figure supplement 1.**
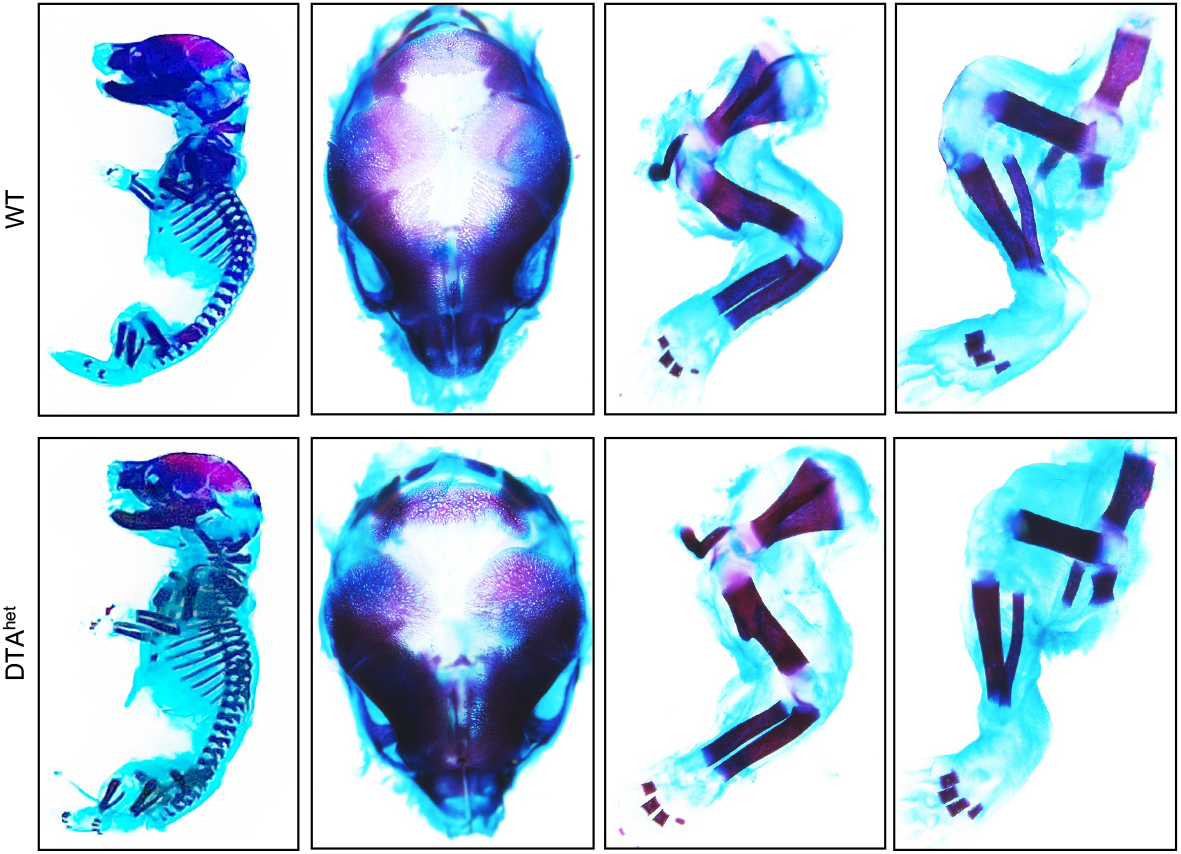
Osteocyte ablation had no impact on embryonic skeletal development. Whole mount skeleton staining of WT and DTA^het^ mice at E19.0 by Alizarin red/Alcian blue.

**Figure 2 - figure supplement 2.**
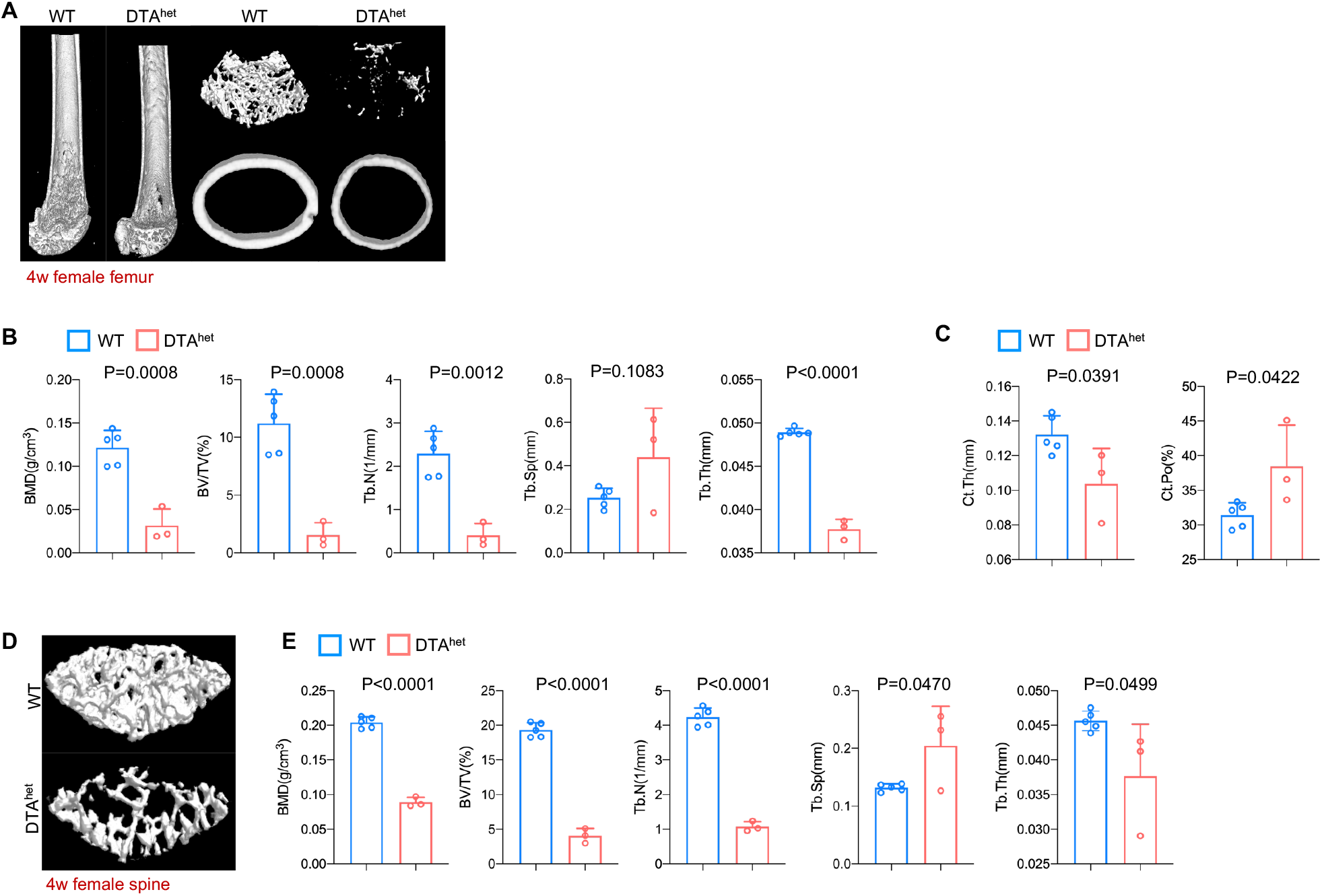
Osteocyte ablation induced severe osteoporosis and kyphosis. A-C Representative μCT reconstructive images of female WT and DTA^het^ mice femur at 4 weeks (A) and trabecular microstructural parameters (BMD, BV/TV, Tb.N, Tb.Sp and Tb.Th) (B) and cortical microstructural parameters (Ct.Th and Ct.Po) (C) derived from μCT analysis (n=3-5 per group). D-E Representative μCT reconstructive images of female WT and DTA^het^ mice third lumbar at 4 weeks (D) and trabecular microstructural parameters (BMD, BV/TV, Tb.N, Tb.Sp and Tb.Th) (E) derived from μCT analysis. Error bar represents the standard deviation.

**Figure 2 - figure supplement 3.**
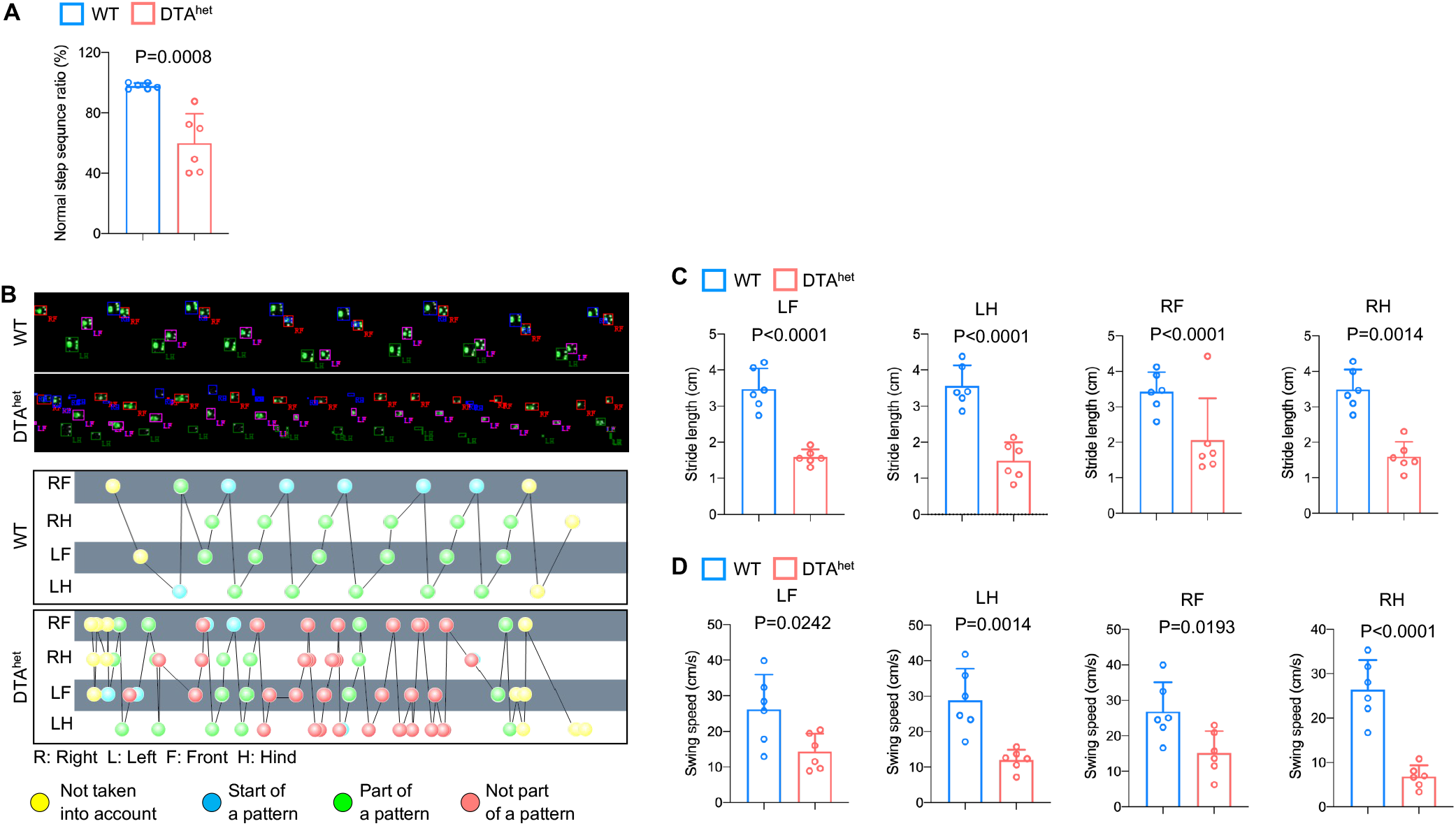
Osteocyte ablation induced severe osteoporosis and kyphosis. A Gait analysis of normal step sequence ratio of male WT and DTA^het^ mice at 4 weeks (n=6 per group). B-D Representative gait images and foot pattern of male WT and DTA^het^ mice (B) at 4 weeks and gait analysis of stride length and swing speed of each paw (C and D). Error bar represents the standard deviation.

**Figure 4 - figure supplement 1.**
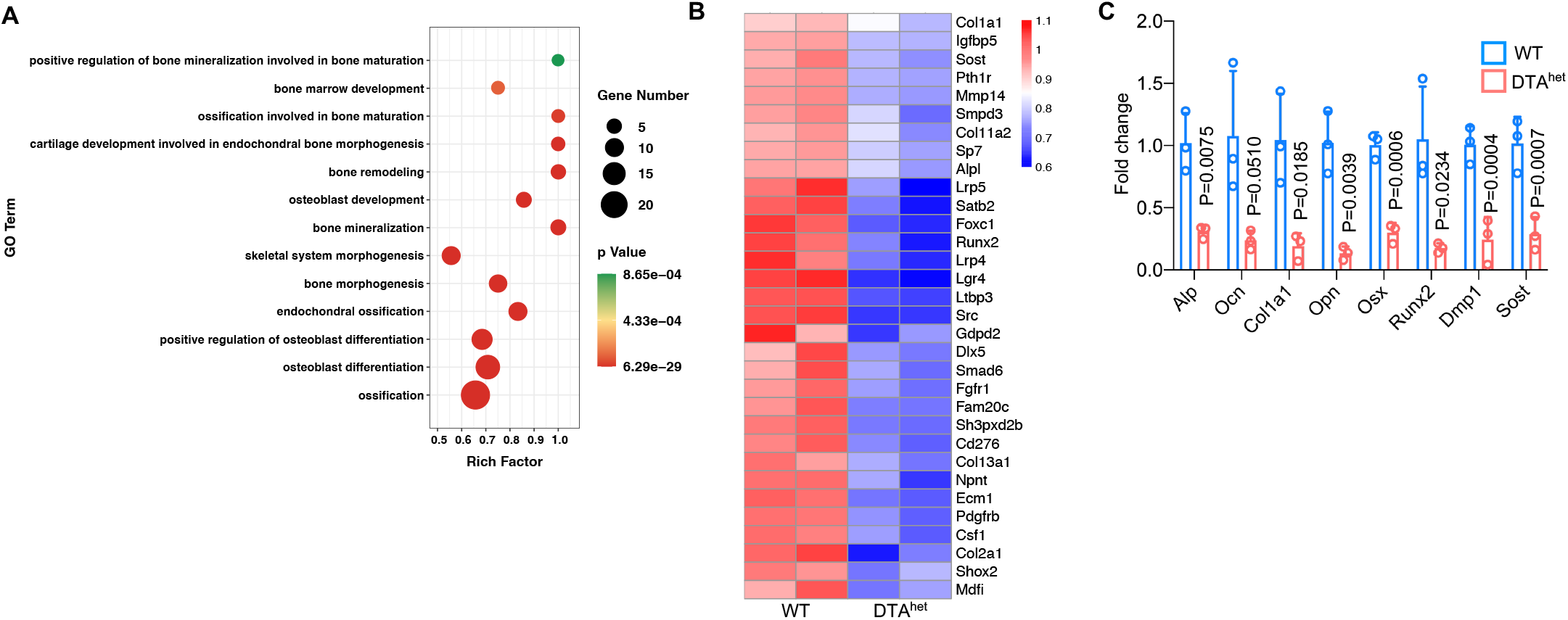
Ablation of osteocytes alters mesenchymal lineage commitment and promoted osteoclastogensis. A Selected osteogenesis related gene ontology (GO) analysis of downregulated genes by osteocyte ablation. B Heatmap of significantly differentiated genes (fold change > 2.0-fold, WT FPKM > 10, FDR < 0.05) (n=2 per group). C Indicated gene expression analysis of the cortical bones of WT and DTA^het^ mice (n=3 per group). Error bar represents the standard deviation.

**Figure 5 - figure supplement 1.**
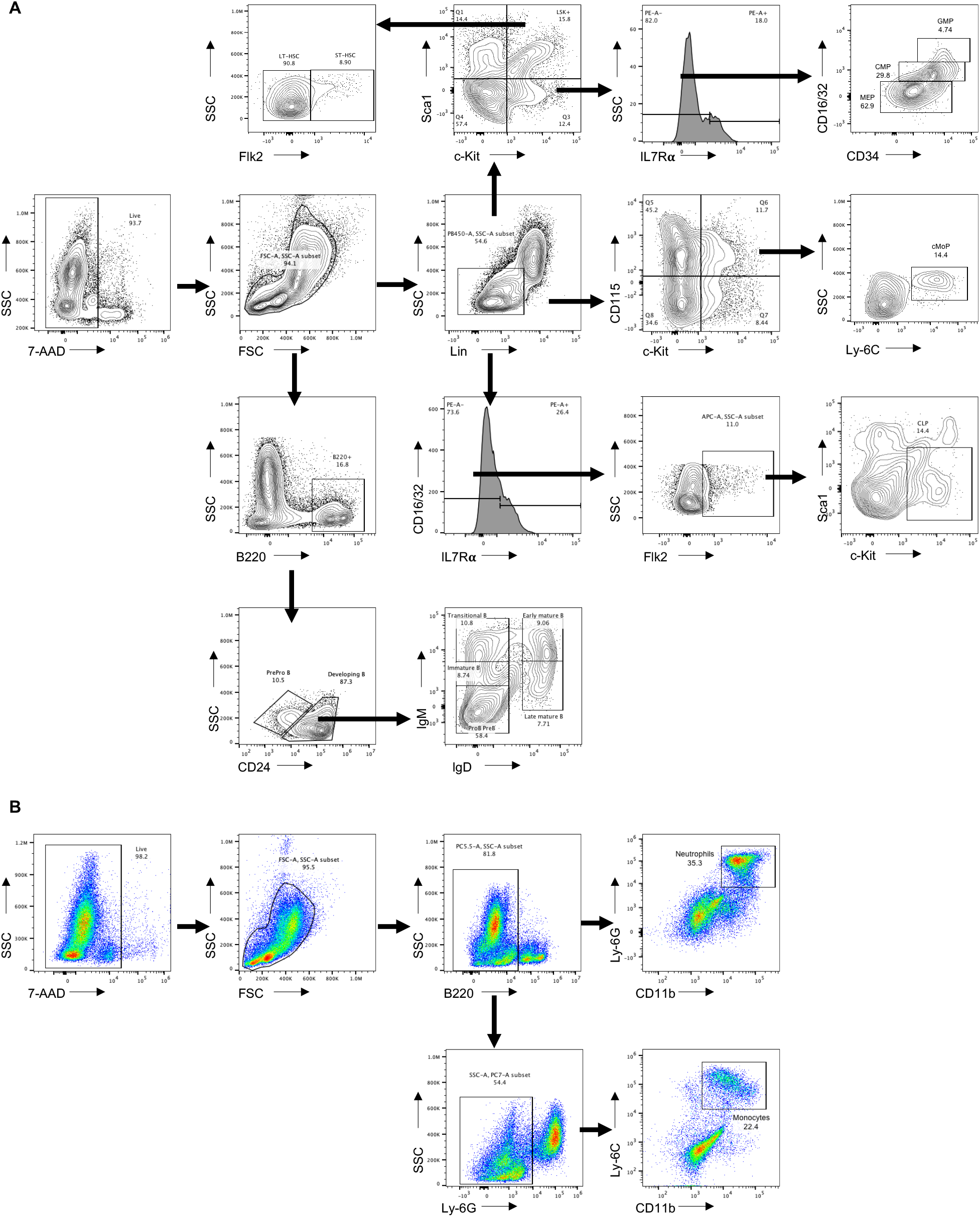
Flow cytometry gating strategy. A Flow cytometry of gating HSC (Lin^-^Sca1^+^c-Kit^+^), LT-HSC (Lin^-^Sca1^+^c-Kit^+^Flk2^-^), ST-HSC (Lin^-^ Sca1^+^c-Kit^+^Flk2^+^), CMP (Lin^-^Sca1^+^c-Kit^-^IL7R*α* ^-^CD34^+^Fc*γ* RII/III^lo^), GMP (Lin^-^ Sca1^+^c-Kit^-^IL7R*α* ^-^CD34^+^Fc*γ* RII/III^hi^), MEP (Lin^-^Sca1^+^c-Kit^-^IL7R*α* ^-^CD34^-^ Fc*γ* RII/III^lo^), cMoP (Lin^-^c-Kit^+^CD115^+^Ly-6C^hi^), CLP (Lin^-^IL7R*α* ^-^Flk2^+^Sca1^+^c-Kit^-^), ProB PreB (B220^+^CD24^+^IgM^-^IgD^-^), immature B (B220^+^CD24^+^IgM^lo^IgD^-^), transitional B (B220^+^CD24^+^IgM^+^IgD^-^), early mature B (B220^+^CD24^+^IgM^+^IgD^+^) and late mature B (B220^+^CD24^+^IgM^lo/-^IgD^+^). B Flow cytometry of gating neutrophil (B220^-^CD11b^+^Ly-6G^+^) and monocyte (B220^-^Ly-6G^-^CD11b^+^Ly-6C^+^).

**Figure 5 - figure supplement 2.**
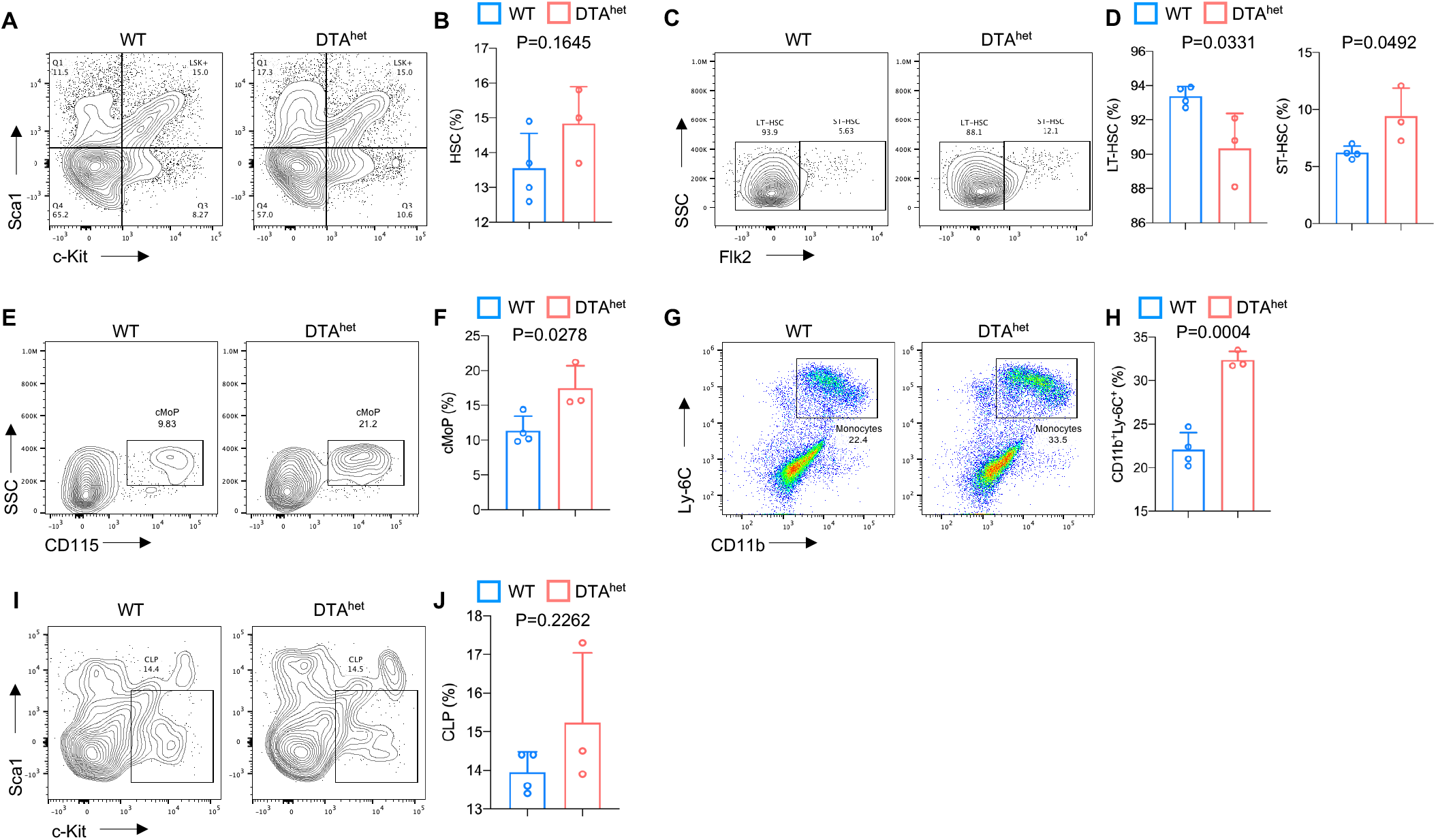
Alteration of hematopoietic lineage commitment by osteocyte ablation. A-B Representative image of flow cytometry (A) and analysis of proportions of HSC (B) of 4 weeks WT and DTA^het^ mice (n=3-4 per group). C-D Representative image of flow cytometry (C) and analysis of proportions of LT-HSC and ST-HSC (D) of 4 weeks WT and DTA^het^ mice (n=3-4 per group). E-F Representative image of flow cytometry (E) and analysis of proportions of cMoP (F) of 4 weeks WT and DTA^het^ mice (n=3-4 per group). G-H Representative image of flow cytometry (G) and analysis of proportions of monocyte (H) of 4 weeks WT and DTA^het^ mice (n=3-4 per group). I-J Representative image of flow cytometry (I) and analysis of proportions of CLP (J) of 4 weeks WT and DTA^het^ mice (n=3-4 per group). Error bar represents the standard deviation.

**Figure 5 - figure supplement 3.**
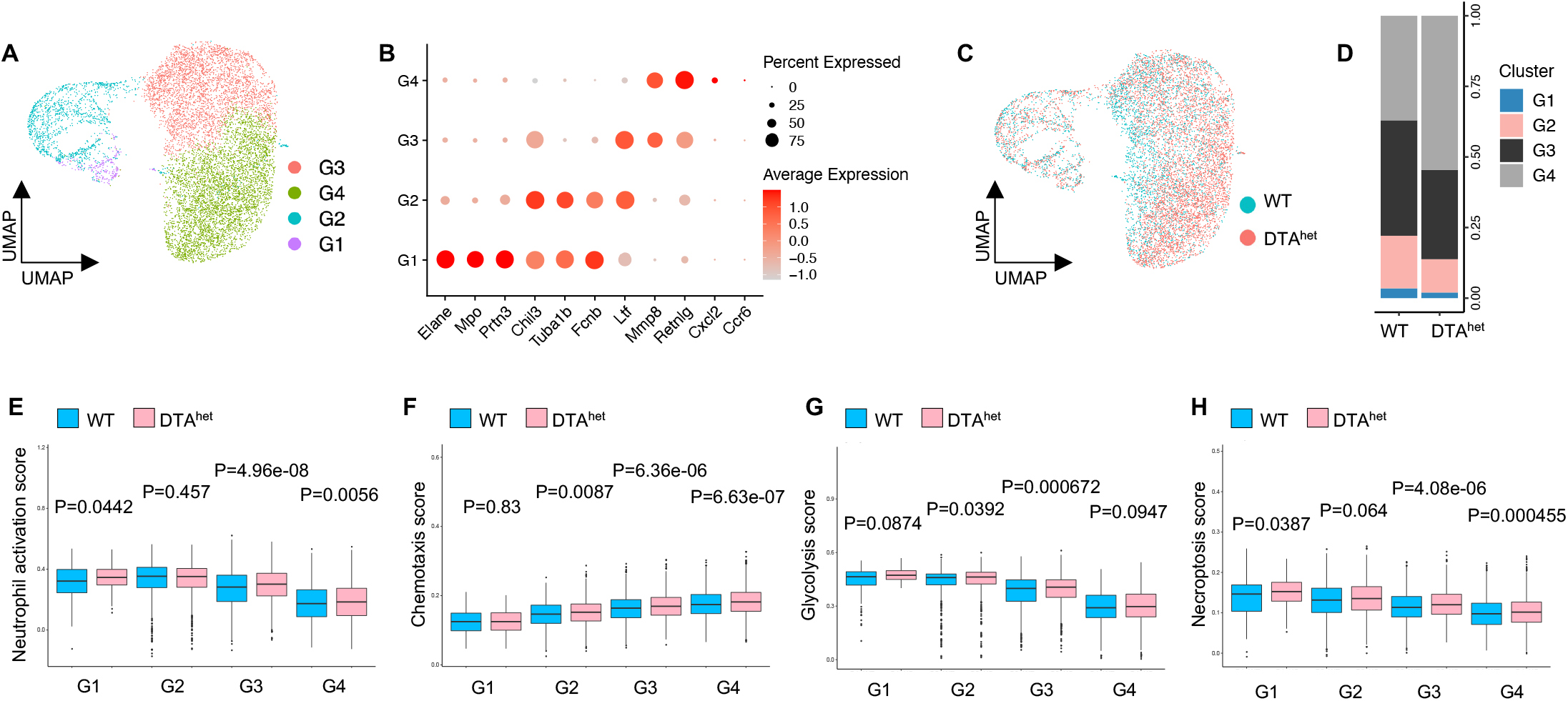
Increased granulopoiesis after osteocyte ablation. A The UMAP plot of neutrophils of 4 weeks WT and DTA^het^ mice and inferred subcluster identity. B Dot plot showing the scaled expression of selected signature genes for each cluster. Dot size represents the percentage of cells in each cluster with more than one read of the corresponding gene and dot are colored by the average expression of each gene in each cluster. C-D The UMAP plot of cells shown by sample (C) and proportions of each subcluster in two samples (D). E-H Comparisons of neutrophil activation score (E), chemotaxis score (F), glycolysis score (G) and necroptosis score (H) between 4 weeks WT and DTA^het^ mice.

**Figure 6 - figure supplement 1.**
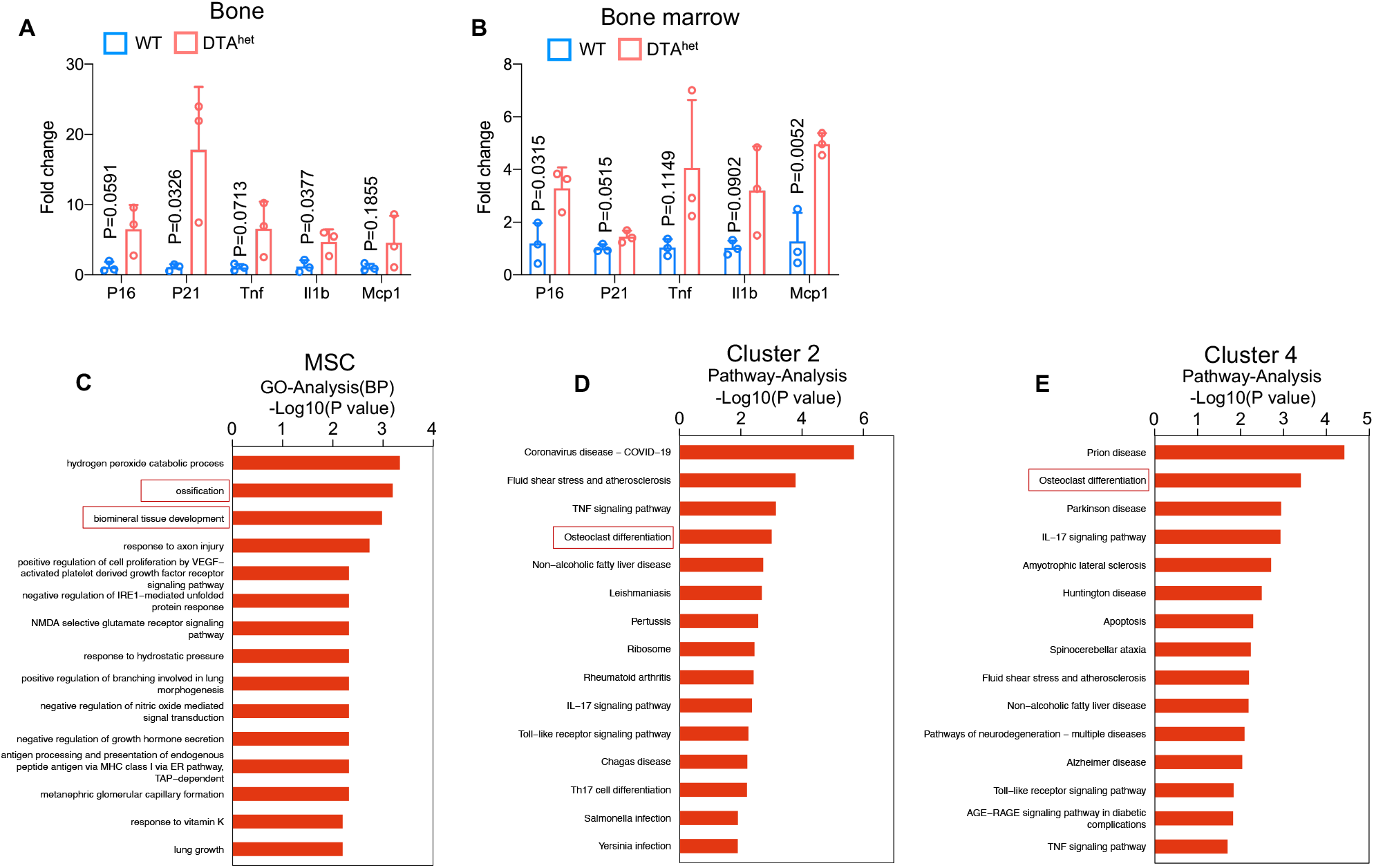
Organismal senescence of osteoprogenitors and myeloid lineage cells leads to the skeletal premature aging. A RT-qPCR analysis of SASP related genes expression at the mRNA level of 4 weeks WT and DTA^het^ mice cortical bone. B RT-qPCR analysis of SASP related genes expression at the mRNA level of 4 weeks WT and DTA^het^ mice bone marrow. C Bar plot of GO analysis of MSC cluster. D-E Bar plot of KEGG analysis of subcluster 2 and 4 of Ly6c2_monocytes. Error bar represents the standard deviation.

